# Deciphering the biology and chemistry of the mutualistic partnership between *Bacillus velezensis* and the arbuscular mycorrhizal fungus *Rhizophagus irregularis*

**DOI:** 10.1101/2023.10.28.564539

**Authors:** Adrien Anckaert, Declerck Stéphane, Laure-Anne Poussart, Stéphanie Lambert, Helmus Catherine, Farah Boubsi, Sebastien Steels, Anthony Argüelles Arias, Maryline Calonne-Salmon, Marc Ongena

## Abstract

Arbuscular mycorrhizal (AM) fungi (e.g. *Rhizophagus irregularis*) recruit specific bacterial species in their hyphosphere. However, the chemical interplay and the mutual benefit of this intricate partnership have not yet been investigated especially as it involves bacteria known as strong producers of antifungal compounds such as *Bacillus velezensis*. Here, we show that the soil dwelling *B. velezensis* migrates along the hyphal network of the AM fungus *R. irregularis*, forming biofilms and inducing metabolic fluxes that contributes to host plant root colonization by the bacterium. During hyphosphere colonization, *R. irregularis* modulates the biosynthesis of specific lipopeptides and antimicrobial compounds in *B. velezensis* as a mechanism toward-off mycoparasitic fungi and bacteria to ensure stable coexistence. These mutual benefits are extended into a tripartite context via the provision of enhanced protection to the host plant through the induction of systemic resistance.

## Main

Arbuscular mycorrhizal (AM) fungi are keystone beneficial microorganisms within rhizobiomes forming symbiotic association with more than two thirds of terrestrial plants. They improve nutrients uptake (P, N) by plants via their extensive and dense extraradical mycelium (ERM) extending tens of cm away from the roots and exploring soil pores via hyphae with diameters up to ten times lower than that of root hairs ^1–3^. The ERM also enables the interconnection of plants of the same or different species to form common mycorrhizal networks (CMNs). Plants can receive warning signals via these CMNs in response to pest and pathogen attack and potentially exchange nutrients^4–7^. Some AM fungal species also have the potential to increase plant systemic resistance (mycorrhiza induced resistance, MIR) against below and above-ground pathogens via mechanisms resembling Induced systemic resistance (ISR)^8–10^.

AM fungi mobilize between 4 and 20% of the total carbon synthesized by plants to feed their own metabolic processes^11^ but also to fuel the catabolism of microbes developing at the hyphal surface in the so-called hyphosphere where carbon rich fungal exudates are released. Like plants do via rhizodeposits, AM fungi thus recruit a wide range of bacterial species forming hyphosphere-associated microbiomes which start to be quite well characterized phylogenetically^12–18^. Some recent studies have highlighted the benefits brought by bacteria to the fungus by the mineralization of organic P and N sources^14,19,20^. Similarly, AM fungal hyphae may serve as conduits for certain phosphate-solubilizing bacteria to reach organic P patches in the soil^21^. However, the molecular basis and phenotypic outcomes of such interkingdom interaction between AM fungi and bacteria dwelling at the hyphal surface is still poorly described^14,18^. In particular, how and to what extent such cooperation could be established with biocontrol bacteria known as strong antagonists remain unexplored, despite the potential benefit of combining these two microorganisms to improve plant protection against pests and diseases.

In this work, we used custom-designed *in vitro* and *in planta* systems with *Solanum lycopersicum* and *Solanum tuberosum* to explore the interkingdom interaction between a well characterized strain of *Rhizophagus irregularis* and *Bacillus velezensis* as model species of plant mutualistic AM fungi and rhizobacteria producing bioactive secondary metabolites (BSMs) with antifungal activity. The *R. irregularis* strain MUCL 41833 was selected for its use in the understanding of AM fungal social interaction with rhizobacteria^19,22^ and its well-known beneficial effects on plant growth and resistance against biotic stresses^23,24^. The GFP-tagged *B. velezensis* strain GA1 was selected for its recognised biocontrol potential and its fully characterized genome and secondary metabolome^25–28^. To study the dynamics of bacterial colonization along the AM fungal ERM, we used a tripartite *in vitro* cultivation system offering the advantage of non-destructive time-lapse observations, allowing the monitoring of interactions between both microorganisms in the absence of any other microbial protagonist. Combining molecular and analytical analyses, we identified key bacterial compounds that contribute to the establishment of a compatible interaction and elucidate how the AM fungus drive the metabolite production in the bacterium and its consequence on antifungal activities of *B. velezensis*. Importantly, we demonstrated that the cooperation between the two microbes confers increased protection of tomato against *Botrytis cinerea*.

## Results

### *B. velezensis* efficiently colonizes the hyphae of *R. irregularis* and expands rapidly along the ERM network

We first studied the spatio-temporal dynamics of the physical interaction between the two microorganisms, using the experimental device schematized in Fig. 1A. Time lapse microscopy imaging revealed that *B. velezensis* cells inoculated at short distance (∼20 µm) from the hyphae of *R. irregularis* moved to and established on the hyphal surface within 48h (Fig. 1B), suggesting a behaviour similar to the chemotaxis seen with plant roots^29,30^. Then, the bacterium spreads over the *R. irregularis* mycelium as motile cells but rapidly transitions to a sessile state, allowing the establishment of multicellular colonies (Supplementary Video 1 and Supplementary Video 2). This leads to the development of a homogenous biofilm along the hyphae as illustrated by the significant increase in colony thickness within the first days post-inoculation (dpi) on short segments (Fig. 1C,D). The *B. velezensis* biofilm rapidly expanded along the runner hyphae (Supplementary Data Fig. 1A), colonizing other structures of the ERM such as spores and branched absorbing structures (BAS) suspected to be devoted to nutrients uptake (Fig. 1E)^31^. Further imaging at a larger scale showed that *B. velezensis* colonized the major part of the dense ERM network (Fig. 1F) 3 dpi (Fig. 1G).

**Fig. 1:**
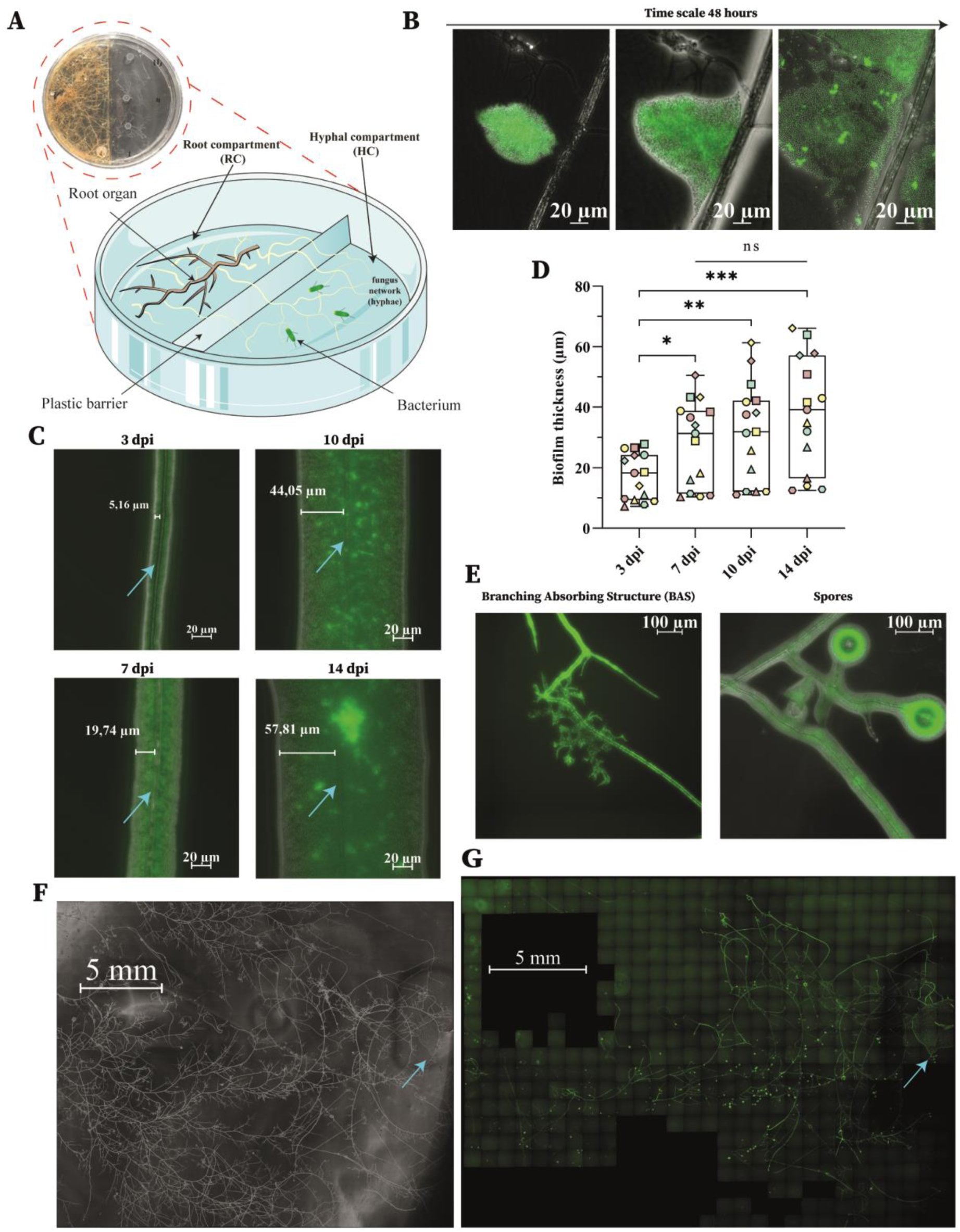
*B. velezensis* behaviour and colonization along AM fungal hyphae. **A**, Picture (red circle) and schematic view of the *in vitro* bi-compartmented Petri plate system allowing to perform live cell microscopy imaging. The Petri plate was separated in two compartments by a plastic barrier, with a root compartment (RC) containing a Ri T-DNA transformed *Daucus carota* root clone DC2 associated to *Rhizophagus irregularis* and a hyphal compartment (HC) containing only hyphae of *R. irregularis*. In the HC, the hyphae were allowed to proliferate on solid MSR without any added carbon source and solidified with agarose (MSR^min^). Once the hyphae were well established in the HC, *Bacillus velezensis* was inoculated on the hyphae to study their interaction. This system offers the advantage of non-destructive time-lapse observations without confounding effects with unwanted microbial contaminants. Brown, transformed root organ of *D. carota*; White, hyphae of *R. irregularis*; Green, cells of *B. velezensis.* B, Microscopic picture of *B. velezensis* GA1 labelled with GFP, attracted and established on the hyphae of *R. irregularis* by chemotaxis (time scale of 48 hours). C, Epifluorescence pictures of *B. velezensis* biofilm development along *R. irregularis.* Epifluorescence pictures were taken by microscopy after 3, 7, 10 and 14 days post-inoculation (dpi) of *B. velezensis* GA1 expressing GFP (green color) on 3-month-old *R. irregularis* hyphae (Blue arrow). By Fiji image processing, measurement of biofilm thickness was achieved on one side of the hyphae. Five co-culture systems were used (biological replicates), and 4 images per co-culture system were taken at different locations over time (technical replicates). Representative pictures are shown in the figures. D, Evolution of *B. velezensis* biofilm thickness along *R. irregularis* hyphae, 3, 7, 10 and 14 dpi. The boxes encompass the 1st and 3rd quartiles, the whiskers extend to the minimum and maximum points, and the midline indicates the median. The individual points represent 5 biological replicates (different shapes) and 3 technical replicates (3 different colours in the same shape). The evolution of the biofilm thickness over time can be followed for each replicate by the same shape and colour. n = 15; one-way analysis of variance (ANOVA) and Tukey’s HSD test (α = 0.05): ns = not significant; * 0.01 < p < 0.05; ** 0.001 < p < 0.01; *** 0.0001 < p < 0.001. E, Epifluorescence pictures of *B. velezensis* biofilm development along *R. irregularis* hyphae and spores. Left: *B. velezensis* development along branched absorbing structures (BAS). Right, *B. velezensis* growth on the surface of spores. Epifluorescence pictures were taken by microscopy respectively 3 and 7 dpi of *B. velezensis* GA1 expressing GFP (green color). F-G, Macroscopic view of the colonization of GFP tagged strain *B. velezensis* GA1 (Green colour in epifluorescence) along the hyphal network of *R. irregularis* 3dpi of *B. velezensis*. Microscopic composite pictures were taken in bright field channel (F) and in epifluorescence (G) The blue arrow show the inoculation drop area on the AM fungal hyphal network.

We calculated an average expansion rate (colonization speed) of 5.6 mm/day on the surface of hyphae, which was significantly higher than observed for the colonization of *Daucus carota* hairy roots (1.2 mm/day) (Fig. 2A and Supplementary Data Fig. 1B). It thus revealed a quite fast invasion of distal parts of the ERM by *B. velezensis* (Fig. 2B) compared to roots (Supplementary Data Fig. 1C). *B. velezensis* colonization density was quantified 3, 7 and 14 dpi by CFU (colony forming units) plate counting. Data showed 20 fold higher populations at the surface of *R. irregularis* hyphae compared with *D. carota* roots (Fig. 2C), further indicating that *B. velezensis* is more prone to colonize *R. irregularis* hyphae than roots in the early stage of interaction. We inferred that this is due to the efficient utilization of fungal exudates by the bacterium leading to a fast grow and formation of an homogenous biofilm along hyphae while root colonization patterns are known to occur with preferential zones of bacterial macrocolony formation^32,33^. However, we also observed that the relative proportion of spores compared to vegetative cells within the *B. velezensis* population colonizing the hyphae of *R. irregularis* was significantly lower (1.6 fold at 7 dpi) than the one colonizing *D. carota* roots in the early days (Supplementary Data Fig. 1D). This high proportion of metabolically active cells in the AM fungal-associated biofilm may thus also reflect a bacterial community more prompt to colonize the fungal host than plant roots.

**Fig. 2:**
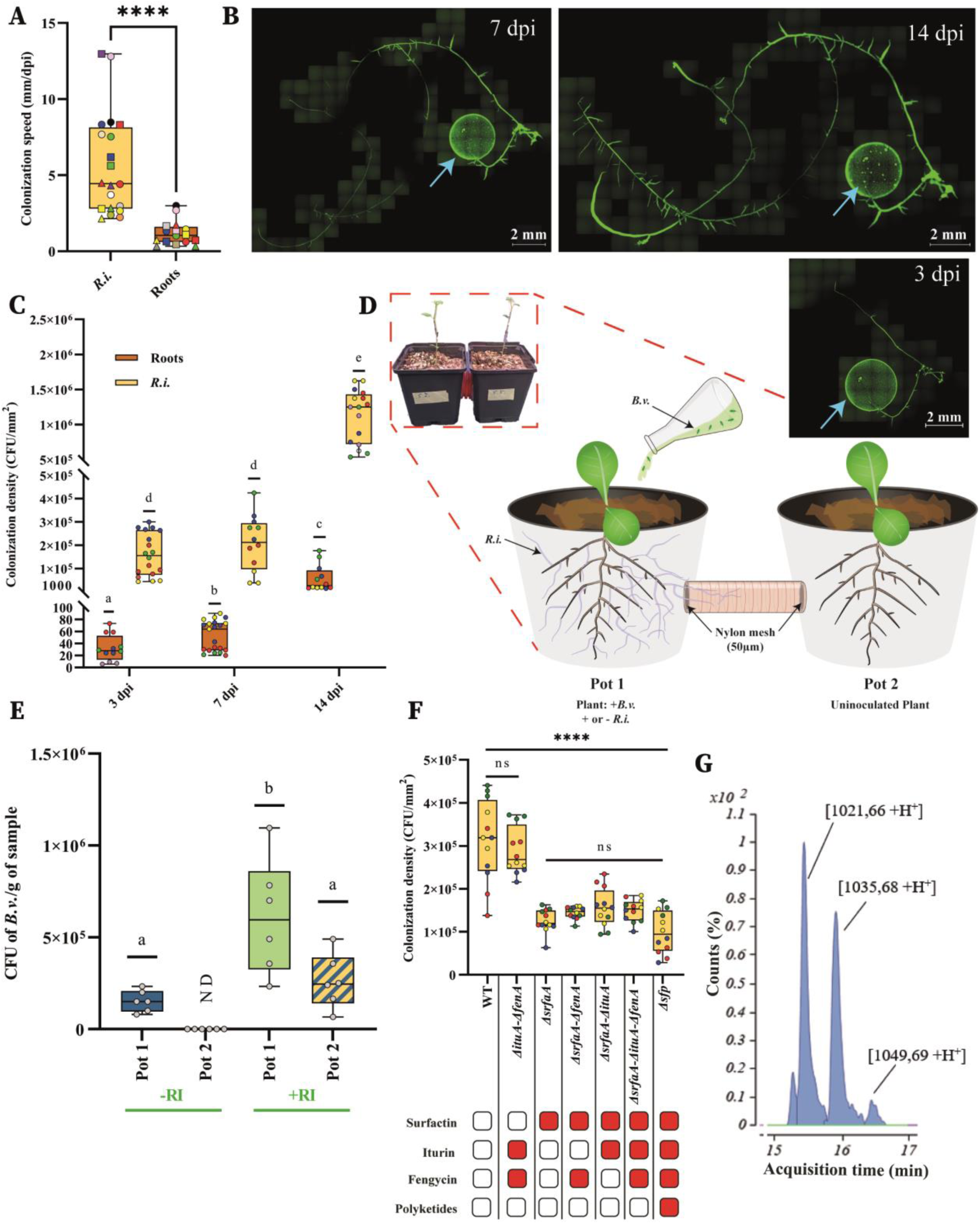
*B. velezensis* colonize effectively AM fungal hyphae compared to root via its surfactin production. **A,** Colonization speed of *B. velezensis* along hyphae of 3-months old cultures of *R. irregularis* (*R.i.* – yellow) and along *D. carota* hairy roots (Roots – brown) during a time lapse of 14 days. The boxes encompass the 1st and 3rd quartiles, the whiskers extend to the minimum and maximum points, and the midline indicates the median. The individual points represent 7 at 11 biological replicates (different colour) and 1-3 technical replicates corresponding to the speed of colonization taken a different time lapse (measured 3 dpi = circle shape, 7 dpi = square shape and 14 dpi = triangle shape). The evolution of the speed of colonization over time can be followed for each replicate with the same colour. 12≤ n ≤ 24; Student’s t-test (α = 0.05): **** = p < 0.0001. **B,** *B. velezensis* colonization along hyphae of 3 month-old cultures of *R. irregularis* overtime. Microscopic composite pictures were taken in epifluorescence. The blue arrow show the inoculation drop area on the AM fungus hyphal network. The speed colonization is quantified by the measurement of the distance travelled by *B. velezensis* (green colour) from the inoculation drop (Blue arrow) divided by the dpi. **C**, Colonization density of hyphae of 3 month-old *R. irregularis* cultures (*R.i.-* yellow) or transformed roots of *D. carota* Cultures (Roots – brown) 3, 7 and 14 dpi of *B. velezensis* The boxes encompass the 1st and 3rd quartiles, the whiskers extend to the minimum and maximum points, and the midline indicates the median. The individual points represent 4 at 6 biological replicates (different colour) and 1 at 6 technical replicates (same colour). 12≤ n ≤ 21; one-way analysis of variance (ANOVA) and Tukey’s HSD test (α = 0.05). Groups with different letters differed significantly from each other at an α of 0.05. **D,** Picture (red square) and schematic view of the experimental design in which two plants of *Solanum tuberosum* (Pot 1 and Pot 2) were connected by a pipe with closed end by nylon mesh of 50 µm porosity. The nylon barrier prevented the passage of roots. During the planting, the Pot 1 was inoculated (+ *R.i*.) or not (*-R.i.*) with *R. irregularis.*. After 5 month of growth, the plants associated or not with *R. irregularis* in Pot 1 were inoculated with *B. velezensis.*. The Plants in Pot 2 were not inoculated by any microorganisms. **E**, Bacterial population of GFP tagged strain *B. velezensis* GA1 quantified by plate-counting of colony forming units (cfu) by gram of sample, 10 dpi. All the plants in pot 1 associated (+*R.i.*) or not (− *R.i.*) with *R. irregularis* Were inoculated with *B. velezensis*. Plants in pot 2 were not inoculated with any microorganisms. The boxes encompass the 1st and 3rd quartiles, the whiskers extend to the minimum and maximum points, and the midline indicates the median. The individual points represent 6 biological replicates (6 different systems). n =6; one-way analysis of variance (ANOVA) and Tukey’s HSD test (α = 0.05). Groups with different letters differed significantly from each other at an α of 0.05. ND = no detected. **F,** Colonization density of hyphae of 3 month-old *R. irregularis* cultures 7 dpi of *B. velezensis* wild type (WT) and mutants unable to produce one or several bioactive secondary metabolites (BSMs). Metabolites not produced by the different mutants are illustrated with red boxes in the table below. The boxes encompass the 1st and 3rd quartiles, the whiskers extend to the minimum and maximum points, and the midline indicates the median. The individual points represent 4 biological replicates (different colour) and 3 technical replicates (same colour). n = 12; one-way analysis of variance (ANOVA) and Tukey’s HSD test (α = 0.05). ns = not significant; **** p < 0.0001**. G**, UPLC-MS extract ion chromatogram (EIC) illustrating the relative abundance of surfactin (blue) secreted by *B. velezensis* 9 dpi on hyphae of a 3-month-old *R. irregularis* culture. The different peaks correspond to the structural variants differing in fatty acid chain length.

We next wanted to test *B. velezensis* colonization under more realistic conditions using mycorrhized *S. tuberosum* plants in greenhouse conditions. We used a homemade setup with two pots, each containing a single plant, connected by a plastic tube covered at both ends with a fine nylon mesh (50 µm diam. porosity) allowing only hyphae of *R. irregularis* to connect the plants (Fig. 2D). In that system, the bacterium inoculated on one mycorrhized plant was able to colonize the plant in the other pot only via migration along the hyphae connecting both plants (Supplementary Data Fig. 1E). Selective plate counting of *B. velezensis* among the bacterial community naturally present in the substrate was based on antibiotic resistance and fluorescence of GFP-tagged GA1. *R. irregularis* was quantified by qPCR using a strain specific house-keeping gene. Data first revealed that plant roots associated with *R. irregularis* (Pot 1 +*R.i.*) exhibited a significantly higher number of GA1 CFUs compared with non-mycorrhized plants (Pot 1-*R.i.*), indicating that the presence of *R. irregularis* favours *B. velezensis* root colonization also in these conditions (Fig. 2E). Moreover, our results showed that *B. velezensis* inoculated on a plant associated with *R. irregularis* (Pot 1 +*R.i.*), was able to colonize a non-bacterized neighbouring plant (Pot 2 +*R.i.*) via the CMN connecting both plants while in the control system without *R. irregularis* (Pot 1-*R.i.*), no bacterial cells were detected on the plant in Pot 2 (Fig. 2E). These findings indicate that the association with *R. irregularis* facilitates the migration of *B. velezensis* cells from one to another root system.

### Surfactin lipopeptide contributes to *B. velezensis.* colonization of hyphosphere

Bacterial fitness can be influenced by secondary metabolites playing key roles in various developmental processes including motility and biofilm formation^34–36^. Thus, a range of mutants specifically repressed in the synthesis of BSMs were tested in order to determine their role in the AM fungal colonization potential of *B. velezensis.* The bacterial population at the surface of hyphae 7dpi was significantly reduced for all the knockout mutants unable to produce surfactin family (Fig. 2F). No other non-ribosomal BSMs were involved in hyphal colonization as indicated by the similar loss in colonization ability observed for the *Δsfp* mutant (repressed in 4′-phosphopantetheinyl transferase which is essential for the proper functioning of non-ribosomal peptide biosynthesis machineries), and other double or triple mutants unable to produce at least surfactin (Fig. 2F). In *B. velezensis* and related species of the *B. subtilis* complex, it has been shown that surfactin deficiency causes impaired biofilm formation, the strains becoming unable to colonize roots^37,38^. Based on these data, we wanted to confirm the production of surfactin by *B. velezensis* upon AM fungal colonization and generated hyphosphere extracts that were analysed by UPLC-qTOF-MS optimized for detection of minimal amounts, quantification and structural characterization of non-ribosomal BSMs. Based on exact mass and retention time compared with standards, we observed substantial amounts of surfactin (as a mixture of homologues that differ in the length of the fatty acid chain, Fig. 2G) at concentrations close to the micromolar range (0.605 µM in average at 9 dpi). This confirmed that *B. velezensis* cells in biofilm associated with *R. irregularis* readily secrete this cyclic lipopeptide (CLiP), which significantly contributes to colonization.

### *B. velezensis* associates with *R. irregularis* in a compatible interaction

*B. velezensis* efficiently colonized the hyphae of *R. irregularis* but this species is a strong producer of antimicrobials and we thus wanted to evaluate the impact of *B. velezensis* on hyphae viability. We first measured succinate dehydrogenase (SDH) activity as an indicator of AM fungal viability via histochemical staining (Fig. 3A)^39^. Image analysis revealed that the enzymatic activity did not differ between non-colonized and colonized hyphae (Fig. 3B) indicating that the presence of the bacterium did not influence adversely the respiratory metabolism of *R. irregularis.* We also monitored via time lapse microscopy imaging, the impact of *B. velezensis* on the cytoplasmic flux within hyphae which is considered as functional trait of AM fungi, allowing the translocation of resources from plant to fungus and vice-versa and somehow reflecting fungal vitality (Supplementary Video 3)^40,41^. Flux velocity inside non-colonized or colonized hyphae remained stable over time, as observed at 3 and 14 dpi. However, the presence of the bacterium increased the cytoplasmic flux velocity compared to non-colonized hyphae whatever the time of observation (Fig. 3C). Thus, *B. velezensis* colonization did not affect fungal fitness, which is in agreement with the fact that the presence of the bacterium does not cause a reduction of the AM fungal population *in planta* (Fig. 3D).

**Fig. 3:**
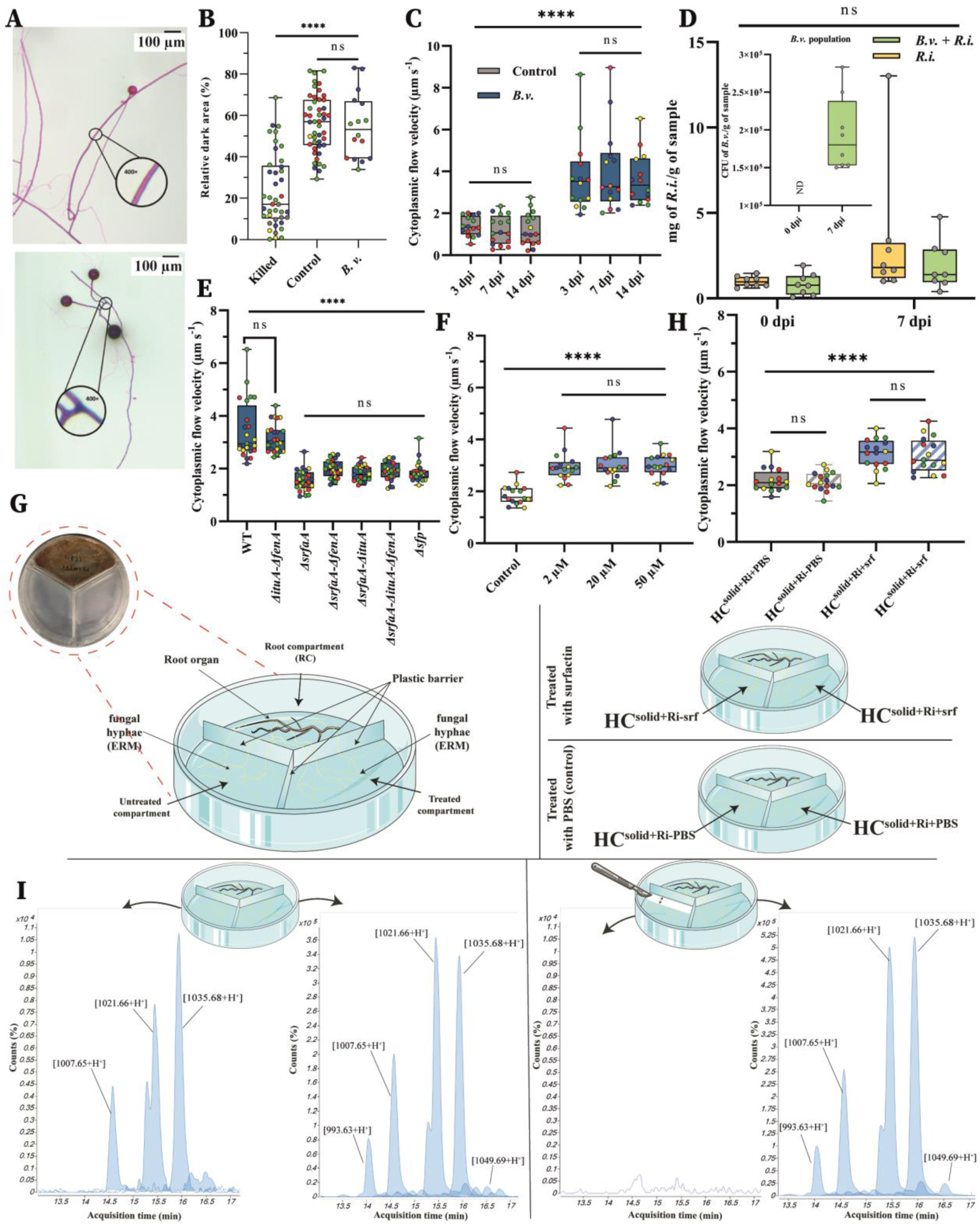
Viability of *R. irregularis* and cytoplasmic streaming within hyphae upon colonization by *B. velezensis*. **A**, Histochemical determination of succinate dehydrogenase (SDH) activity indicating respiratory metabolism of *R. irregularis* hyphae. (**Bottom**) Control (with physiological water) showing dark blue-violet zones corresponding to SDH activity. (**Top**) killed hyphae (with formaldehyde) showing pink color corresponding to dead hyphae. Pictures of stained hyphae from a 3-month old culture of *R. irregularis* were taken with the stereomicroscope. The images presented are representative examples selected from independent samples repeated on minimum 3 biological replicates. **B,** Relative dark area corresponds to potential SDH activity of *R. irregularis.* colonized by *B. velezensis* (*B.v.*) 14 dpi along hyphae of 3-month old cultures of *R. irregularis*, compared to killed hyphae (with formaldehyde 2%) and the control (treated with physiological water). The boxes encompass the 1st and 3rd quartiles, the whiskers extend to the minimum and maximum points, and the midline indicates the median. The individual points represent 3 to 5 biological replicates (different colours) and 1 to 20 technical replicates (same colours). 16 ≤ n ≤ 47; one-way analysis of variance (ANOVA) and Tukey’s HSD test (α = 0.05): ns = not significant; **** = p < 0.0001. **C,** Cytoplasmic flow velocity in hyphae of *R. irregularis* in presence or absence of *B. velezensis* tagged GFP. measured 3, 7 and 14 dpi. The boxes encompass the 1st and 3rd quartiles, the whiskers extend to the minimum and maximum points, and the midline indicates the median. The individual points represent 4 to 8 biological replicates (different colours) and 1 to 6 technical replicates (same colours). 14 ≤ n ≤ 24; one-way analysis of variance (ANOVA) and Tukey’s HSD test (α = 0.05): ns = not significant; **** p < 0.0001**. D,** Fungal population of *R. irregularis* (*R.i.*) quantified by quantitative PCR reported by gram of roots sampled with attached substrate (mg of *R.i*. /g of sample), 0 and 7 dpi of plants treated with inoculum of *B. velezensis* (*B.v.+R.i*., green) or with solution with any microorganisms (*R.i.*, yellow). The boxes encompass the 1st and 3rd quartiles, the whiskers extend to the minimum and maximum points, and the midline indicates the median. The individual points represent 8 biological replicates (8 different plants). n =8; one-way analysis of variance (ANOVA) and Tukey’s HSD test (α = 0.05): ns = not significant. Inside the graph is included the bacterial population of *B. velezensis* in the treatment combining *B. velezensis* and *R. irregularis* (*B.v.+R.i*., green), before the inoculation of the bacterium and 7 dpi of *B. velezensis*. **E,** Cytoplasmic flow velocity in hyphae of *R. irregularis*. in presence of *B.velezensis* wild type (WT) or knock out mutants, 7 dpi along hyphae. The boxes encompass the 1st and 3rd quartiles, the whiskers extend to the minimum and maximum points, and the midline indicates the median. The individual points represent 4 biological replicates (different colours) and 6 to 7 technical replicates (same colours). 24 ≤ n ≤ 25; one-way analysis of variance (ANOVA) and Tukey’s HSD test (α = 0.05): ns = not significant; **** p < 0.0001**. F,** Cytoplasmic flow velocity in hyphae of *R. irregularis* in contact with increasing concentrations of pure surfactin (2µM, 20µM or 50 µM) compared to a PBS control. The boxes encompass the 1st and 3rd quartiles, the whiskers extend to the minimum and maximum points, and the midline indicates the median. The individual points represent 4 biological replicates (different colours) and 4 technical replicates (same colours). n=16; one-way analysis of variance (ANOVA) and Tukey’s HSD test (α = 0.05): ns = not significant; **** p < 0.0001**. G,** Picture (red circle) and illustration of a tri-compartmented experimental setup. In one compartment (root compartment – RC) roots of *D. carota* are associated to *R. irregularis*, from which the hyphae colonize two side compartments. One side compartment contained either pure surfactin at a concentration of 2µM (HC^solid+Ri+srf^) or PBS (HC^solid+Ri+PBS^). The other side compartment received no surfactin or PBS was annotated as follows (HC^solid+Ri-srf^ or HC^solid+Ri-PBS^). A plastic barrier prevent surfactin from diffusing between compartments. **H,** Cytoplasmic flow velocity in the compartment treated with surfactin (HC^solid+Ri+srf^, blue) or PBS (HC^solid+Ri+PBS^, gray) and in the untreated compartment following the treated compartment either with surfactin (HC^solid+Ri-srf^, blue stripes) or with PBS (HC^solid+Ri-PBS^, gray stripes). The boxes encompass the 1st and 3rd quartiles, the whiskers extend to the minimum and maximum points, and the midline indicates the median. The individual points represent 4 biological replicates (different colours) and 4 technical replicates (same colours). n=16; one-way analysis of variance (ANOVA) and Tukey’s HSD test (α = 0.05): ns = not significant; **** p < 0.0001**. I,** Illustration of the tri-compartmented experimental setup in which surfactin was added to one hyphal compartment (Treated compartment), keeping the same amount of surfactin 19.16 µM in average (n=3) than the concentration used to the treatment (20µM). In the second compartment, the amount of surfactin was quantified when the ERM was intact (left) or when the network was cut in a band 0.5 cm width (right) to evaluate if the surfactin may be translocate by *R. irregularis*. Representative UPLC-MS extract ion chromatogram (EIC) illustrating the relative abundance of surfactin (blue) shown for each compartment. *D. carota* transformed roots were confined to the root compartment (RC), but the fungus was able to cross the plastic barrier, entering the hyphal compartments (HC). Another plastic barrier prevented the transfer of surfactin between the two HC.

Some secondary metabolites and more particularly CLiPs are key components involved in multitrophic interactions established by *Bacillus* in the rhizosphere ^25,42^. We thus investigated their possible role in the increase of cytoplasmic flux velocity triggered by the bacterium in *R. irregularis* hyphae. We first analysed GA1 mutants deleted in CLiP-biosynthesis genes. Co-cultivation with *R. irregularis* of all mutants impaired in surfactin production did not result in increased flux velocity, providing a first strong evidence for the crucial role of this CLiP but not iturin, fengycin or any other non-ribosomal product (Fig. 3E). As an important proof we added purified surfactin to monocultures of *R. irregularis* associated to *D. carota* and observed a similar trend on flux velocity upon treatment with concentrations as low as 2 µM, which is in the range of amounts detected in the hyphosphere (Fig. 3F). These data strongly suggest that surfactin serves as signal molecule produced by *B. velezensis* and is perceived by the AM fungus to boost its cytoplasmic translocation. In order to determine if this response is restricted to hyphae segments in contact with the lipopeptide or not, we next used a tri-compartmented culture system where the hyphal network but not the root, is allowed to cross into two different hyphal compartments physically separated by a plastic barrier, thereby preventing surfactin from diffusing between the compartments (Fig. 3G). With this set-up, we observed the diffusion of the response within the ERM and across the root compartment. The effects of surfactin were not limited to hyphae in the treated compartment (HC^solid+Ri+srf^) but also influenced distal regions (Fig. 3H), resulting in an overall increase in cytoplasmic flux observed in the untreated compartment (HC^solid+Ri-srf^) at 48 hpi (Fig. 3H and Supplementary Data Fig. 2). These data strongly suggest that surfactin serves as signal molecule produced by *B. velezensis* and is perceived by the AM fungus to systemically boost its cytoplasmic translocation. Interestingly, UPLC-qTOF MS analyses of hyphosphere extracts prepared from the untreated compartment (HC^solid+Ri-srf^) revealed, that surfactin was translocated at distal zone through the ERM (Fig. 3I) since significant amounts (corresponding to 0.32 µM in average) were recovered from the HC^solid+Ri-srf^ when the ERM was intact but no trace of the lipopeptide could be detected when the mycelium network was cut. These results underlined the potential of the AM fungus to transport this bacterial secondary metabolite through its network.

### Attenuated fengycin production in the hyphosphere prevents *B. velezensis* from antagonizing *R. irregularis*

*B. velezensis* is a strong producer of antifungal BSMs including iturin– and fengycin-type CLiPs which are well described for their activity against a wide range of fungal plant pathogens^25,43,44^. In order to understand how *R. irregularis* may co-exist with this antagonistic bacterium, we tested the toxicity of pure CLiPs on the fungus. We used propidium iodide (PI) staining as indicator of membrane integrity since toxicity of these molecules mainly relies on their pore-forming activity in biological membranes causing cytosolic leakage and death of target cells ^45–47^. Fengycin and iturin did not impact hyphae membrane integrity at 2 µM, while at 20 and 50 µM, fengycin markedly destabilized the membranes and iturin only at the highest concentration (50 µM) (Fig. 4A,B). In contrast, surfactin did not affect membrane integrity over the tested concentration range (Fig. 4B).

**Fig. 4:**
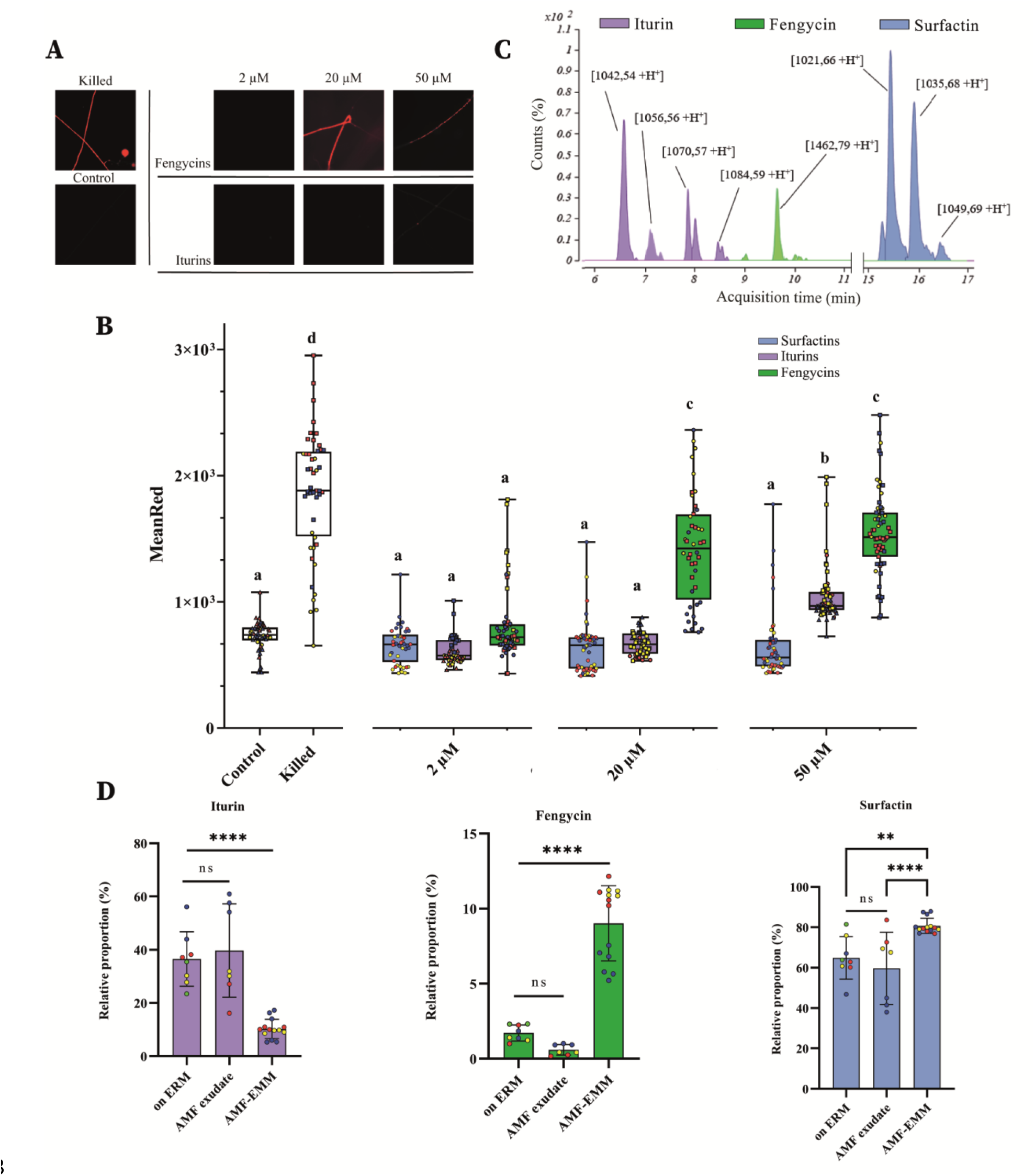
Effect of *B. velezensis* antifungal compounds on *R. irregularis* and their modulation through *R. irregularis* exudates. **A-B**, *R. irregularis* hyphae stained with 50µg ml^-1^ Propidium Iodide after treatment with 2µM, 20µM or 50µM of fengycin, iturin or surfactin. AM fungal hyphae treated with PBS solution (Control) and 1% Triton X-100 with 2% formaldehyde (Killed) were used as controls. A, Propidium iodide cell staining of *R.i.* hyphae observed by fluorescence microscopy. B, Dose effect of pure fengycin (green), iturin (purple) and surfactin (blue) produced by *B. velezensis* on membrane integrity of hyphae of 3 month old (triangle shape), 5 month old (square shape) or 6 month old (circle shape) *R. irregularis* cultures measured by fluorescence upon staining with propidium Iodide. High MeanRed values mean high membrane destabilization. The boxes encompass the 1st and 3rd quartiles, the whiskers extend to the minimum and maximum points, and the midline indicates the median. The individual points represent 3 biological replicates (different colours) and 9 to 26 technical replicates (same colours). The dose effect of fengycin, iturin and surfactin have been evaluated according to different ages of *R.irregularis* cultures (different shapes). 43≤ n ≤ 65; Letters a to d indicate statistically significant differences according to one-way analysis of variance (ANOVA) and Tukey’s HSD test (α = 0.05). C, UPLC-MS extract ion chromatogram (EIC) illustrating the relative abundance of surfactin (blue), iturin (purple) and fengycin (green) family secreted by *B. velezensis* after 9 dpi on hyphae of 3 month old cultures of *R.irregularis.* The different peaks for each CLiPs correspond to the structural variants differing in fatty acid chain length. D, Relative surfactin (blue), iturin (purple), fengycin (green) proportion corresponding to the detected peaks areas of each CLiP compared to the total amount of the 3 families of CLiPs following growth condition. CLiPs relative proportion when *B. velezensis* evolved along AM fungal hyphae (on ERM), on exudates collected from AM fungal hyphae (AM fungal exudate), on AM fungal exudates mimicking medium (AMF-EMM). Bars represent the means ± SD of 3 to 4 biological replicates (different colours) and 2 to 4 technical replicates (same colours). 7 ≤ n ≤ 14; Letters a to d indicate statistically significant differences according to one-way analysis of variance (ANOVA) and Tukey’s HSD test (α = 0.05): ns = not significant; ** 0.001 < p < 0.01, **** p < 0.0001.

Next, we wanted to evaluate the production of these compounds in the hyphosphere upon AM fungal colonization. CLiP profiling via UPLC-qTOF-MS revealed a distinct pattern compared to the one observed upon growth in lab media or in a medium mimicking plant exudates^48^ (Fig. 4C). Upon growth in rich optimized lab media, *B. velezensis* typically secretes CLiPs in relative proportions of approx. 50% surfactin, 25% iturin and 25% fengycin ^48^. However, the hyphosphere samples exhibited different ratios, with surfactin accounting for 60% (± 11.21%) and iturin for 36.5% (± 10.20%), while fengycin was present in minimal amounts of only 1.5% (± 0.78%)(Fig. 4D). The amounts recovered corresponded approximately to 0.61 µM ± 0.21 for surfactin, 0.48 µM ± 0.016 for iturin and 0.03 µM ± 0.001 for fengycin. Similar relative CLiPs proportions were observed when *B. velezensis* planktonic cells were cultured in presence of exudates collected from AM fungal hyphae as sole nutrient source (surfactin: 2.73 µM ± 1.04; iturin: 1.86 µM ± 1.06; fengycin: 0.0478 µM ± 0.0325), indicating that the reduced fengycin production is not related to biofilm formation and is not contact-dependent (Fig. 4D). Since the nature of carbon sources in the medium may influence BSMs production by *Bacillus* spp.^48–50^, we postulated that modulation of the CLiP patterns in the hyphosphere could be driven by the specific nutritional context offered by the hyphae exudates. To simulate this context, we developed a minimal medium called AMF-EMM (AM fungus exudate mimicking medium) that only contains oligo-elements and the carbon sources typically encountered in hyphal exudates^13–15^. Cultivating *B*. *velezensis i*n this medium resulted in a significant increase in the relative proportions of fengycin and surfactin compared with natural exudates (Fig. 4D). Therefore, we assume that the CLiP pattern produced by *B. velezensis* in interaction with *R. irregularis* and characterized by very low amounts of harmful fengycin is not due to specific nutritional context but is formed in response to the perception of some unidentified signal(s) secreted by the AM fungus.

### *B. velezensis* produces antimicrobials inhibiting soilborne competitors

Our data show that the *B. velezensis* cell population evolving as biofilm on AM fungal hyphae still efficiently produces bioactive secondary metabolites and we wanted to know if this could provide some protection to its fungal host against potentially harmful microbes such as *Trichoderma harzianum* and *Collimonas fungivorans*^51,52^. We first evaluated the growth inhibitory activity of the crude cell free supernatant (CFS) obtained after growth of *B. velezensis* in AM fungal exudates towards *T. harzianum* and *C. fungivorans* and observed a significant antagonist effect of the CFS from wild-type GA1 on both species (Fig. 5A; Fig. 5B). We next tested mutants of *B. velezensis* unable to produce those BSMs that are readily formed by GA1 wild-type upon growth in the hyphosphere. A complete loss of anti-*Trichoderma* activity was observed by testing CFS extracts obtained from the mutants *ΔbacA* and *ΔsfpΔbacA* repressed in the synthesis of the non-ribosomal SFP-independent di-peptide bacilysin (Fig. 5A). The crucial role of bacilysin in the antifungal activity developed by *B. velezensis* against *T. harzianum* was further supported by the similar activity of the *Δsfp* mutant unable to form the other non-ribosomal products CLiPs and polyketides (Fig. 5A). Significant amounts of bacilysin were detected in hyphosphere extracts (Fig. 5C) indicating that this compound was readily formed by *B. velezensis* upon AM fungal colonization. Results with mutants repressed in the synthesis of antifungal CLiPs known for their individual or synergistic antifungal activity, demonstrated the reduced role of these metabolites against *T. harzianum* within the hyphosphere context (Supplementary Data Fig. 3). Previous research has highlighted the significance of iturin production by *B. velezensis* in inhibiting the growth of *Trichoderma* spp ^53^. However, we assume that in our conditions, the concentration of iturin in the hyphosphere did not reach the level necessary to affect the development of this mycoparasite. Regarding *C. fungivorans*, none of the knockout mutants showed a significant loss in the antibacterial activity observed for the wild type (Fig. 5B and Supplementary Data Fig. 3B). Conserved anti-*Collimonas* activity in Δ*sfp* and Δ*sfp*Δ*bacA* extracts suggests the involvement of additional ribosomal compound(s) that remains to be identified.

**Fig. 5:**
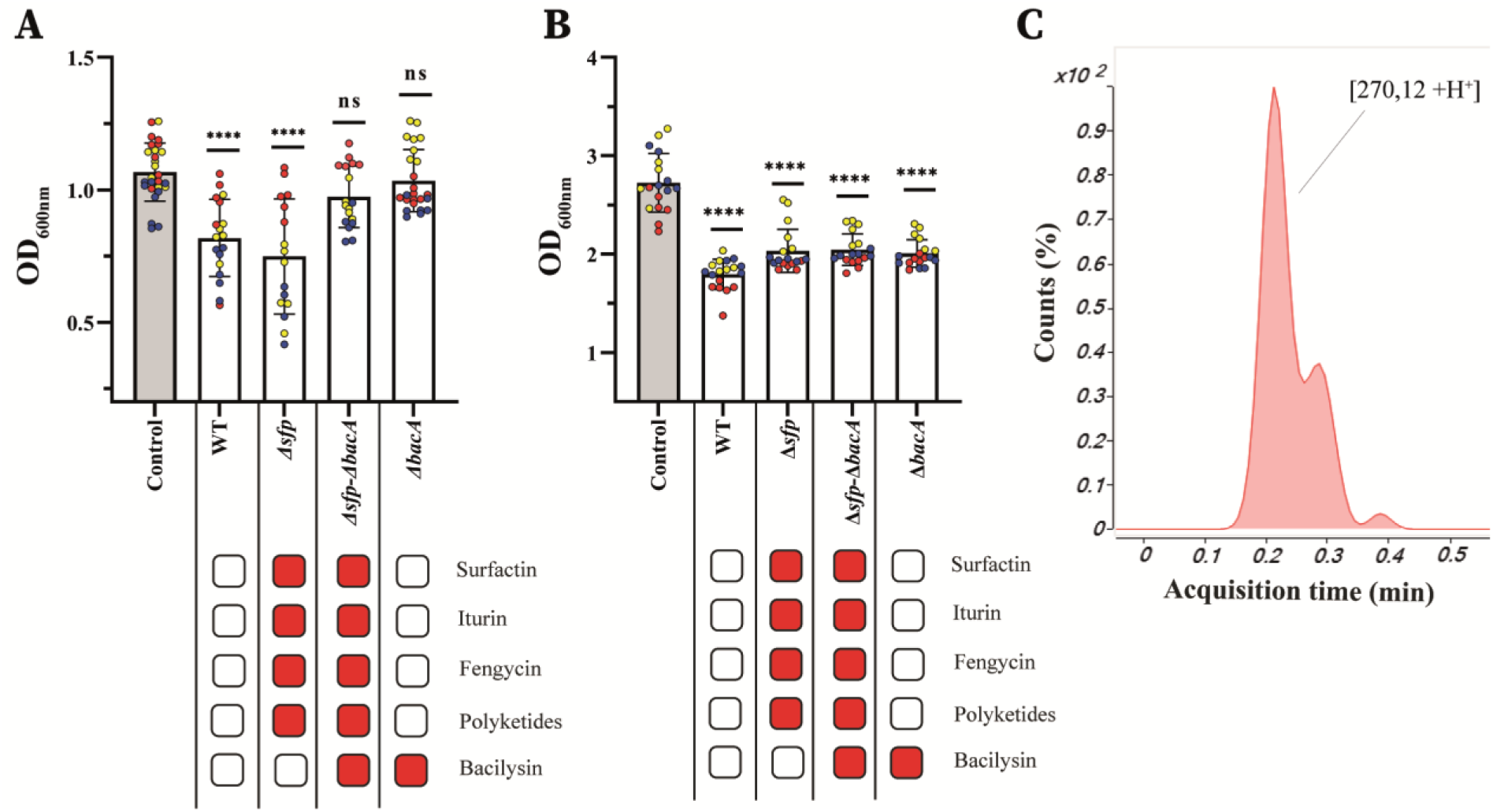
BSMs produced by *B. velezensis* in the hyphosphere of *R.irregularis* allowing an antagonism activity against *Trichoderma harzianum* and *Collimonas fungivorans*. **A**, Effect of GA1 wild type or mutant cell-free supernatants (CFS), produced on exudates of *R.i.* cultures, on the growth of *Trichoderma harzianum* Rifai MUCL 29707. The optical density (OD_600nm_) of the fungi was quantified after 36h of growth in presence or in absence (control) of CFS. Metabolites not produced by the different mutants are illustrated with red boxes in the table below. Bars represent the means ± SD of 3 biological replicates (different colours) and 4 to 9 technical replicates (same colours). 16 ≤ n ≤ 27, one-way analysis of variance (ANOVA) and Dunnett’s test (α = 0.05): ns = not significant; **** = p < 0.0001. B, Effect of GA1 wild type or mutant cell-free supernatants (CFS), produced on exudates of *R.i.* cultures on the growth of *Collimonas fungivorans* LMG 21973. The optical density (OD_600nm_) of the bacterium is quantified after 24h of growth in presence or in absence (control) of CFS. Metabolites not produced by the different mutants are illustrated with red boxes in the table below. Bars represent the means ± SD of 3 biological replicates (different colours) and 6 technical replicates (same colours). 16 ≤ n ≤ 27, one-way analysis of variance (ANOVA) and Dunnett’s test (α = 0.05) compared to the control: ns = not significant; **** = p < 0.0001. C, UPLC-MS extract ion chromatogram (EIC) illustrating the relative abundance of bacilysin (red), secreted by *B.v.* 9 dpi on 3 month old *R.i*. cultures.

### *B. velezensis* and *R. irregularis* interaction provides enhanced ISR functionality

In a more applied perspective for biocontrol application, we next tested in greenhouse trials whether the combination of *B. velezensis* and *R. irregularis* is able to protect tomato plants (*Solanum Lycopersicon*, used as model for *Solanaceae*) from infection with *Botrytis cinerea,* a major pathogen in a range of crops^54,55^. In the system used, we assessed the protective effect due to systemic resistance induced by the beneficial microbes inoculated at the root level against disease caused by *B. cinerea* on leaves in the form of spreading necrotic lesions. Data showed a strong decrease in both disease severity (approx. 60%) and disease incidence (approx. 75%) in plants co-inoculated with *R. irregularis* and *B. velezensis* compared to controls. Treatments with the AM fungus or the bacterium alone provided some protection but to a significantly lower level (Fig. 6A,B). Microbial population monitoring in soil of co-inoculated plants (as described in Fig. 2D) revealed that the AM fungal density remained stable overtime while *B. velezensis* CFUs markedly increased within two weeks (Fig. 6C,D), confirming first that the bacterium does not negatively impact the fungus and second that the presence of the AM fungus may favour bacterial soil invasion. Moreover, inoculation with both microorganisms did not impact plant size compared to controls (Fig. 6E), strongly suggesting that enhanced disease resistance is not indirectly due to a higher robustness. From all these data, we infer that mutualistic cooperation between *R. irregularis* and *B. velezensis* confers an increased potential for immune activation in tomato plants and higher systemic resistance.

**Fig. 6:**
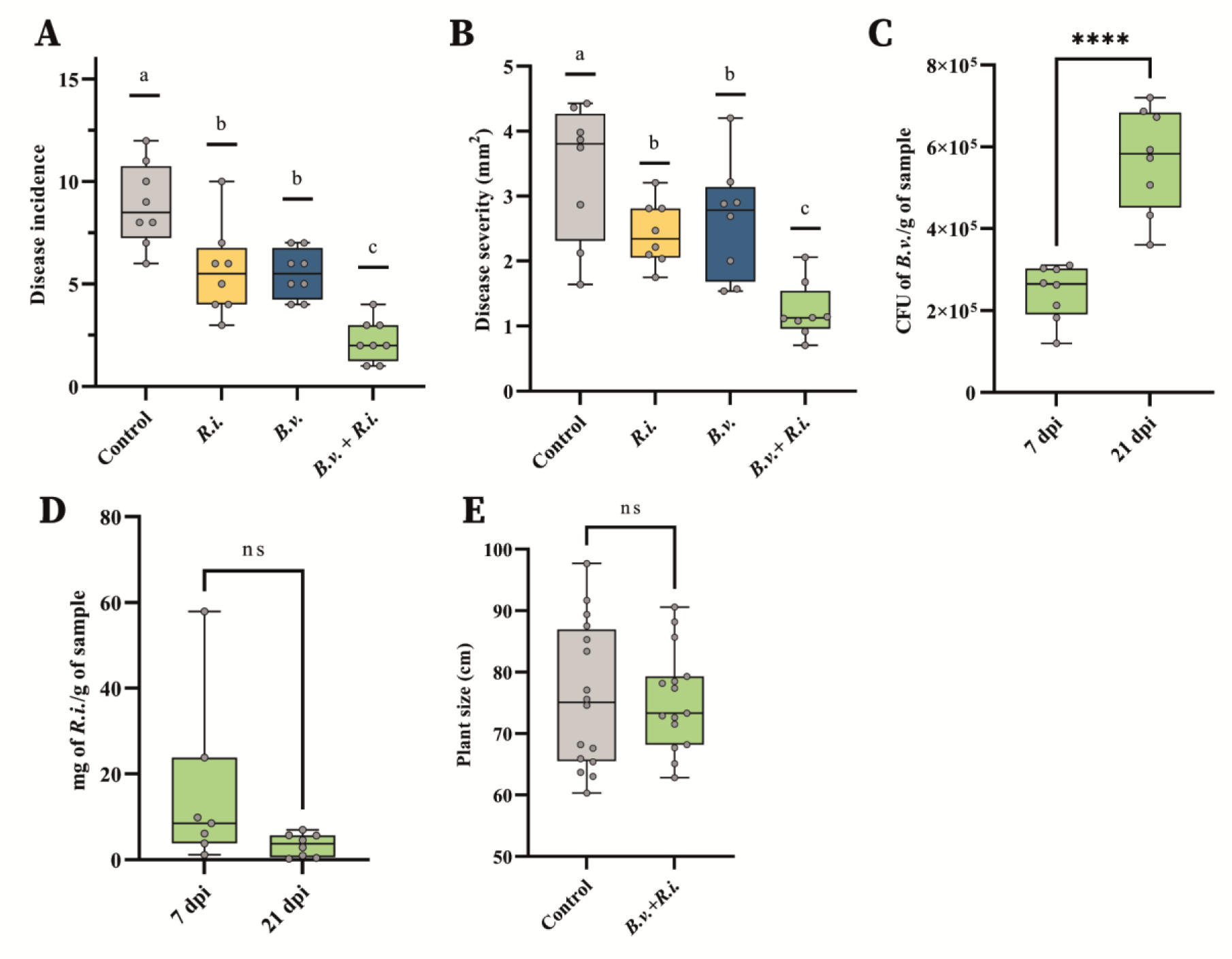
Interaction between *B. velezensis* and *R.irregularis* increases plant protection. **A**, Number of emerging/spreading lesions– (Disease incidence) 21 days after infection with *B. cinerea* evaluated on 15 leaflets of 8 *S. lycopersicon* plants treated with *R. irregularis* (*R.i.*, yellow)*, B. velezensis* (*B.v.,* blue) or the combination (*R.i.+ B.v*., green) or with Hoagland solution as Control (gray). The boxes encompass the 1st and 3rd quartiles, the whiskers extend to the minimum and maximum points, and the midline indicates the median. The individual points represent 8 biological replicates. n=8; Letters a to c indicate statistically significant differences according to one-way analysis of variance (ANOVA) and Tukey’s HSD test (α = 0.05). **B**, Size of emerging lesions (Disease severity) quantified by ImageJ Fiji, 21 days after infection with *B. cinerea*. on leaves of *S. lycopersicon* plants treated with *R. irregularis* (*R.i.*)*, B. velezensis* (*B.v.*) or with Hoagland solution as Control. The boxes encompass the 1st and 3rd quartiles, the whiskers extend to the minimum and maximum points, and the midline indicates the median. The individual points represent 8 biological replicates. n=8; Letters a to c indicate statistically significant differences according to one-way analysis of variance (ANOVA) and Tukey’s HSD test (α = 0.05). **C,** Bacterial population of GFP tagged strain *B. velezensis* GA1 quantified by plate-counting of colony forming units (CFU) reported by gram of roots sampled with attached substrate (CFU of *B.v.*/g of sample), 7 and 21 dpi of the bacterium on *S. lycopersicon* roots associated with *R. irregularis* (*B.v.+R.i*., green). The boxes encompass the 1st and 3rd quartiles, the whiskers extend to the minimum and maximum points, and the midline indicates the median. The individual points represent 8 biological replicates. n =8; Student’s t-test (α = 0.05): **** = p < 0.0001. **D,** Fungal population of *R. irregularis* quantified by qPCR reported by gram of roots sampled with attached substrate (mg of *R.i*. /g of sample), 7 and 21 dpi of the bacterium on the *S. lycopersicon* plants treated with the combination *R. irregularis* and *B. velezensis* (*B.v.+R.i*., green). The boxes encompass the 1st and 3rd quartiles, the whiskers extend to the minimum and maximum points, and the midline indicates the median. The individual points represent 8 biological replicates. n =8; Student’s t-test (α = 0.05): ns = not significant. **E,** Tomato plant size treated with *R. irregularis* associated with *B. velezensis,* 7 dpi of the bacterium (before treatment with *B. cinerea*) compared with the control treated with Hoagland solution. The boxes encompass the 1st and 3rd quartiles, the whiskers extend to the minimum and maximum points, and the midline indicates the median. The individual points represent 16 biological replicates. n =16; Student’s t-test (α = 0.05): ns = not significant.

## Discussion

In most instances, cross-kingdom interactions between plant beneficial rhizobacteria and soilborne fungi result in antagonistic outcomes and fungal growth inhibition due to production of secondary metabolites with fungicidal activity^25,56,57^. Here we unveil a rather unanticipated compatibility between *B. velezensis* as strong bacterial competitor and *R. irregularis* as keystone AM fungal species, ensuring a stable coexistence and partnership. *B. velezensis* dwelling in the hyphosphere efficiently produces iturin-type lipopeptides known for their strong antifungal properties against a wide range of phytopathogens^43,58–60^ but which appears not toxic to *R. irregularis* at the concentrations tested (2-20 µM). In general, the biological activity of lipopeptides is mainly related to their ability to interact with the cell membrane of the target organism, which depends both on the structure of the molecule and on the lipid composition and organization of the target membrane^45–47^. AM fungal membranes, are mainly constituted of 24-methyl cholesterol and phosphatidylinositol/phosphatidylcholine with saturated and/or monounsaturated fatty acids as phospholipids and this widely differs from lipid compositions in pathogenic fungi ^61–63^. Therefore, we assume that the low toxicity of iturin to *R. irregularis* is due to the specific lipid content within the plasma membrane of AM fungal hyphae. In contrast to iturin, our data reveal that fengycin exhibit antagonistic activity at low micromolar concentration on AM fungal hyphae such as reported for many phytopathogenic fungi^44,64,65^. However, production of this CLiP by *B. velezensis* in the hyphosphere is very low and the compound does not accumulate at inhibitory amounts in the vicinity of AM fungal hyphae. Based on our results, we postulate that fengycin synthesis is dampened in response to the perception of some signal emitted by *R. irregularis.* Beyond their role as nutrients supporting bacterial growth, some carbohydrates and carboxylates secreted by the AM fungi have been described as signals triggering phenotypic and metabolic responses in bacteria^19,66^. However, this is obviously not the case here since growing *B. velezensis* in artificially reconstituted medium containing the typical sugars, organic acids and amino acids exuded by *R. irregularis* does not lead to such fengycin repression. Other AM fungal secreted products may putatively function as signals like effector proteins/peptides or plant-derived metabolites such as methyl salicylate known to be involved in plant-microbe cross-talk but none of these compounds could be detected upon UPLC-MS analysis of exudates collected from *R. irregularis* cultures^4,7,67,68^. Resolving the chemical nature of the *R. irregularis* signalling molecule(s) responsible for fengycin modulation thus deserves further investigation but still, in a broader context, our observation paves the way to the discovery of AM fungal compounds driving cross-kingdom interactions. Our understanding of the external biotic signals that modulate the synthesis of bioactive secondary metabolites in *Bacillus* spp. is still limited ^25,44^, and this study highlights AM fungi as unique soil-dwelling microbes that may impact CLiP production.

Our findings also provide insights into the molecular basis driving mutualism between the two microbes. The lipopeptide surfactin plays a key role as it not only contributes to efficient hyphosphere invasion (by favouring motility and biofilm formation), as reported for root colonization^27,37,38^, but also acts as a signal sensed by the fungus for boosting its cytoplasmic flux and hence its vitality and functionality. This represents a new natural function for this lipopeptide as signal mediating interkingdom interactions even if deciphering the mechanistic of surfactin perception by the AM fungus requires further investigation. However, based on the similarity of membrane lipids between AM fungi and plants, we hypothesize that it may rely on a specific interaction with the lipid phase of the AM fungal plasma membrane by analogy with what is observed for interactions with plant cells in the context of immunity stimulation^27,69–71^. Interestingly, surfactin in its canonical form is ubiquitously synthesized by all species belonging to the *B. subtilis* group that are widespread in soil. Other structurally-related CLiPs are produced by other rhizobacteria such as *Pseudomonas*^72,73^. This suggests that this type of cross-kingdom communication mediated by CLiPs may have a more global distribution belowground.

Our *in vitro* data further indicate that *B. velezensis* readily uses hyphal exudates to eavesdrop on the AM fungus and fuel its catabolism to sustain growth as described for other species ^15,66^, but also to promptly form biofilm and invade the hyphosphere in a process more efficient than rhizosphere colonization. Our observations need to be broadened to other AM fungal species^14^ but *B. velezensis* seems to behave as a true fungiphile perfectly adapted to live in association with AM fungi. Thought they are not considered as members of the core hyphosphere microbiome, some bacilli have been recurrently identified as dominant components within the bacterial community associated with AM fungi^16,74–78^. This suggests that some species like *B. velezensis* underwent specific adaptation during evolution allowing it to thrive in a lifestyle compatible with AM fungi.

Efficient colonization of ERM provides clear ecological advantages to the bacterium. First, biofilm formation is an essential trait that protects the cell community against abiotic stresses, as well as against infiltration by competitors or toxins via the shield effect of the hydrophobin layer^29,79,80^. Secondly, by using the large and dense hyphal networks of AM fungi as dynamic support for biofilm establishment and expansion, *B. velezensis* may extend in a volume of soil inaccessible to roots and thus considerably enhance its invasiveness and persistence in the niche. Bacterial migration along AM fungal hyphae over significant distances has been only recently demonstrated for the phosphate solubilizing bacteria *Rhanella aquatilis*^21^. Here we provide first strong evidence on spatio-temporal dynamics of hyphal invasion by *B. velezensis* extending much further from the inoculation zone as previously observed in other fungal-bacterial interaction ^14,21,66^. In support to the mutualistic nature of the interaction, our findings highlight the functional significance of *B. velezensis* related to the fitness of AM fungi. The biofilm formed by *B. velezensis* may also serve as protective shield for its AM fungal host as illustrated by the toxicity of the secretome of *B. velezensis* dwelling in the hyphosphere toward potentially harmful microorganisms such as *T. harzianum* and *C. fungivorans*. Although we could not identify the full range of predicted BSMs, *B. velezensis* in biofilm efficiently synthesizes multiple antimicrobial compounds that not only protect cells in biofilm against invasive organisms but that may also form a chemical barrier that contributes in safeguarding the AM fungi from microbial aggressors to compensate for the low natural potential of AM fungi to produce antibiotic weapons. We thus infer that AM fungi may selectively recruit members of the hyphosphere, such as *B. velezensis,* as protective agents, thereby expanding functionalities of the AM fungal-associated microbiome beyond their role in facilitating nutrient uptake^14,18^. This broadens the concept of the hyphosphere’s impact on the selection of specific functional groups of bacteria in the hyphosphere of AM fungi to include antagonistic and biocontrol species such as *B. velezensis*.

We also show that the interaction between *R. irregularis* and *B. velezensis* positively impacts host plant health via enhanced protection against *Botrytis cinerea* infection since combination of the two microorganisms confers higher systemic resistance compared with their application as single bioinoculants. On the one hand, this may be due to an enhanced MIR functionality of the AM fungus triggered by the bacterium but nothing is known about the nature of fungal elicitors or effectors that could be boosted upon interaction. On the other hand, surfactin is the main compound produced by *B. velezensis* acting as elicitor of immune responses and systemic resistance in tomato and other Solanaceae^27,33,69,71^ and we hypothesize that its translocation via the AM fungal ERM network may provide an optimal delivery at the root level in high amounts. A higher accumulation of surfactin at the root surface could also result from a higher population of *B. velezensis* since soil invasion by the bacterium is clearly facilitated in presence of the AM fungus. *Bacillus spp.* used as monospecies bioinoculants do not always meet expectations in the level and consistency of disease protection provided to crops, which is mainly due to poor or insufficient establishment of threshold populations in the soil environment under natural conditions once introduced. Assessing whether such synergy between *R. irregularis* and *B. velezensis* may help combat other diseases via ISR or direct antagonism requires further investigations but combination of the two microorganisms seems very promising as a new type of microbial consortium to be implemented in agricultural systems for sustainable crop production. A better understanding of the nature and dynamics of cross-kingdom interactions between these two microorganisms provides a way forward to engineering consortia with predictable compatibility and high biocontrol potential.

## Materials and methods

### Biological materials

*Rhizophagus irregularis* (Błaszk,Wubet, Renker, and Buscot) C. Walker and A. Schüßler as (“irregulare”) MUCL 41833 was obtained from the Glomeromycota *in vitro* collection (GINCO). The fungus was proliferated *in vitro* on Ri T-DNA transformed roots of carrot (*Daucus carota* L.) clone DC2 in bi-compartmented Petri plates (90 × 15 mm) containing the Modified Strullu-Romand (MSR) medium.^81^ The Petri plates were incubated in the dark at 27 °C until sufficient spores were produced.

*Bacillus velezensis* GA1 and its mutants are listed in Supplementary Table 1. The construction of knockout mutant strains of *B. velezensis* GA1 involved gene replacement through homologous recombination as previously described by Hoff *et al.*^27^. The procedure involved PCR amplification of the 1 kb upstream region of the target gene, the antibiotic marker (chloramphenicol, kanamycin and/or phleomycin cassette), and the downstream region of the target gene, using appropriate primers. The primers used for this study have been previously described by Andric *et al.*^26^. To introduce the recombinant cassette into GA1, a slightly modified protocol based on the method by Jarmer *et al.*^82^ was employed by inducing natural competence through nitrogen depletion. Initially, a colony of *B. velezensis* was inoculated on lysogeny broth (LB) medium (10 g l^−1^ NaCl, 5 g l^−1^ yeast extract, and 10 g l^−1^ tryptone) and incubated at 37°C under shaking for 6 hours. The cells were then washed and resuspended. Subsequently, the recombinant cassette (1 µg) was added to the GA1 cell suspension, adjusted to an OD_600_ of 0.01. The incubation was carried out at 37°C under shaking for 24 hours. Colonies that had integrated the cassette through a double crossing over event were selected on LB plates supplemented with chloramphenicol (5 µg ml^-1^), phleomycin (4 µg ml^-1^) or kanamycin (5µg ml^-1^). The successful gene deletions were confirmed by PCR analysis using specific upstream and downstream primers (UpF and DwR) and by the absence of production of the corresponding bioactive secondary metabolites. Mutants of *B. velezensis* were grown on LB medium supplemented with adapted antibiotics.

*Trichoderma harzianum* Rifai MUCL 29707 was obtained from the Mycothèque de l’Université catholique de Louvain (BCCM/MUCL). The strain was reactivated and periodically cultured on PDA. *Collimonas fungivorans* LMG 21973 was grown on TSA medium (Sigma-Aldrich, India) and obtain from Microbial Processes and Interactions (MiPI) laboratory.

*Solanum tuberosum* L. var. Bintje was provided by the “Station de Haute Belgique” (Libramont, Belgium) as *in vitro* plants. The plants were micropropagated every 4 weeks in culture microboxes, sealed with breathing filters in the lid (ref: 0118/120 + OD118, SacO_2_, Belgium). They were grown on sterilized (121°C for 15 min) Murashige and Skoog (MS) (Duchefa, Netherlands) medium supplemented with 10 g l^−1^ sucrose and solidified with 4.2 g l^−1^ phytagel (Sigma-Aldrich, St. Louis, USA). The microboxes were placed in a growth chamber (Snijders Scientific B.V., Netherlands), under a temperature of 20/18°C (day/night), a relative humidity (RH) of 75%, a photoperiod of 16 h day^−1^ and a photosynthetic photon flux (PPF) of 50 μmol s^−1^ m^−2^.

*Solanum lycopersicum* cv Ailsa Criag and cv MoneyMakers seeds were obtained from EnGraineToi (Sussargues, France). Seeds underwent sterilization by immersion in 70% (v/v) ethanol for 1 min and 10% (v/v) sodium hypochlorite (NaOCl, 12%) for 4 min, then thoroughly rinsed in sterile water before their used in experiments.

The protective effect of tomato due to systemic resistance induced was performed using *Botrytis cinerea* MUCL 43839 as pathogen agent. The strain was obtained from BCCM/MUCL.

### Set up of experimental *in-vitro* system

Bi-compartmented Petri plates (90 × 15 mm) were used to grow transformed carrot roots with AM fungus as detailed in St-Arnaud et al.^83^. In one compartment (the root compartment – RC), the root and AM fungus were associated on 25 ml MSR medium. The MSR medium was solidified with 4.2 g l^−1^ Phytagel, while in the other compartment (the hyphal compartment – HC), only the extraradical mycelium of the fungus was allowed to grow. The fungus extended in the HC via a slope (from top to bottom of the plastic barrier separating the RC from the HC) made of 5 ml MSR medium, without sucrose and vitamin sources (MSR^min^), gelified with 10 g l^−1^ agarose. Ten ml of the same solid medium was poured in the HC (HC^solid^). After circa 3 months of growth, the extraradical mycelium (ERM) network of *R. irregularis* extended profusely in the HC (thereafter HC^solid+Ri^) (Supplementary Figure 5). A control treatment consisting of non-mycorrhizal excised transformed carrot roots was included and roots of *D. carota* were allowed to cross the partition wall separating RC from HC to grow for circa 3 months in the HC (thereafter HC^solid+Dc^) (Supplementary Figure 5). The system was further developed with the HC containing liquid MSR medium, with the exception of the slope^15^. Briefly, in the HC, 15 ml of liquid MSR^min^ medium diluted twice (MSR^min-½^) was added (HC^liquid^). After 3 months, a profuse ERM has developed in the HC (HC^liquid+Ri^) (Supplementary Figure 5). Bi-compartmented Petri plates were incubated in the dark at 27 °C.

Similarly, to the above, bi-compartmented Petri plates were used to grown *Solanum tuberosum* var Bintje plantlets. Twenty days old *in vitro S. tuberosum* plantlets were transferred in the Petri plates with their shoot protruding outside the plate via a small opening in the lid, plastered with sterile silicon grease to avoid contaminations, and the roots developed in the RC on 20 ml MSR^min^ medium. The roots crossed the partition wall and developed in the HC on the MSR^min^medium gelified with 10 g l^−1^ of agarose (HC^solid+St^)(Supplementary Figure 5). Fresh medium was added weekly to keep medium at the top of the partition wall in the RC. The plants were kept for ∼3 months in a growth chamber (Snijders Scientific B.V., Netherlands), under a temperature of 20/18°C (day/night), a RH of 75%, a photoperiod of 16 h day^−1^ and a photoperiod under a PPF of 50 μmol s^−1^ m^−2^.

In complement to the bi-compartmented Petri plates, a tri-compartmented *in vitro* culture setup was developed. In this system, the roots were paired with the AM fungus in one compartment (referred to as the root compartment – RC). From this RC, the ERM extended into two adjacent compartments (referred to as the hyphal compartments – HC). The compartment containing the root culture was supplied with MSR, whereas the two hyphal compartments lacked sucrose, and vitamin resources (MSR^min^) and gelified with agarose (HC^solid^).In these HCs, only the ERM of the fungus was allowed to grow (HC^solid+Ri^), while the roots were confined to the RC (Fig. 3G).

### Root and hyphae colonization by *B. velezensis in vitro*

After 4 weeks of growth, the ERM in the HC^solid+Ri^ or transformed roots in the HC^Solid+Dc^ treatments were inoculated with the *B. velezensis* tagged GFP and colonization of hyphae or roots monitored by microscopy. For this, the bacterial cells were precultured overnight in liquid root exudates mimicking exudates of *Solanaceae* (RE)^48^ medium with shaking (180 rpm). The cells were then washed twice with physiological water (NaCl 0.9%, w/v) and bacterial concentration adjusted to 7.5×10^8^ CFU ml^−1^. A single *R. irregularis* hyphae or root of *D. carota* was inoculated with one drop of 1 µl of bacterial suspension. The colonization of GA1 tagged GFP was monitored at regular intervals. Microscopic composite pictures were performed within epifluorescence to determinate the speed of bacterial colonization along hyphae or roots of *D. carota*. The speed of colonization was quantified by measuring the distance travelled by *B. velezensis* from the inoculation drop divided by the day post inoculation (dpi). The quantification was performed from the biological material of minimum 7 plates (biological replicates) at 3^rd^, 7^th^and 14^th^ dpi.

Microscopy imaging was performed using a Nikon Ti2-E inverted microscope (Nikon, Japan) equipped with ×20/0.45 NA S Plan Fluor objective lenses (Nikon, Switzerland) and a Nikon DS-Qi2 monochrome microscope camera. Images and videos taken in the bright field channel were acquired using a Ti2 Illuminator-DIA and an exposure time of 20 ms. *B.velezensis* tagged GFP was visualized by conventional epifluorescence microscopy. A lumencor sola illuminator (Lumencor, USA) was used as the source of excitation with an exposure time of 500 ms and the GFP-B HC Bright-Line Basic Filter was used.

### Velocity of cytoplasmic flow

The velocity of cytoplasmic flow inside hyphae of *R. irregularis,* colonized or not by *B. velezensis,* was measured 3, 7 and 14 dpi of the bacterium or for the control, by capturing videos in the bright field channel as described in section “Root and hyphae colonization by *B. velezensis in vitro*”. The inoculation of hyphae by *B. velezensis* or GA1 mutants was also performed as previously described in section “ Root and hyphae colonization by *B. velezensis in vitro*”. The control treatment involved treating non-colonized hyphae with 1 µl of physiological water. The velocity of cytoplasmic flow colonized by *B. velezensis* mutants was evaluated only at 7 dpi.

The cytoplasmic flow velocity of *R. irregularis* was also quantified in the presence of pure surfactin. Hyphae of *R.irregularis* were inoculated with 1 µl of pure surfactin solubilized in PBS at 2 µM, 20 µM or 50 µM concentration. A control treatment with only 1 µl PBS was included.

To investigate the impact of surfactin on the distal regions of the *R. irregularis* network, away from direct contact with the lipopeptide, a second experiment was conducted using a three-compartment Petri plate system. One of the two hyphal compartments was treated (the Treated compartment), receiving either a pure surfactin solution at 2µM (HC^solid+Ri+srf^) or PBS (HC^solid+Ri+PBS^). The second hyphal compartment remained untreated but was labelled according to its proximity to the treated compartment, either with surfactin (HC^solid+Ri-srf^) or with PBS (HC^solid+Ri-PBS^). These three compartments were separated by plastic barriers, preventing direct physical connections between them. However, the fungal network itself remained continuous and interconnected through the root compartment. The velocity of cytoplasmic flow in *R. irregularis* was quantified in all 4 treatment conditions, in each HC.

For all experiments, measurements were conducted on minimum of 4 plates (biological replicates) in which minimum 1 video was taken (technical replicate) following the treatment. The velocity was quantified using Fiji by manual tracking with Trackmate a Fiji plugin^84,85^.

### Succinate dehydrogenase (SDH) activity in hyphae

Histochemical staining was performed to quantify the succinate dehydrogenase (SDH) activity according to the adapted procedures of Schaffer and Peterson^39^. The SDH activity of *R. irregularis* hyphae was assessed 7 and 14 dpi with *B. velezensis* (HC^Solid+Ri^) and compared to hyphae non-inoculated with the bacterium (1 µl of physiological water – 0.9 % (w/v)) and hyphae killed with formaldehyde (2 % (v/v))^86^. The inoculation of hyphae by *B. velezensis* was performed as previously described in section “ Root and hyphae colonization by *B. velezensis in vitro*”. The AM fungal hyphae developing on the surface of the MSR^min^ medium in the HC^Solid+Ri^ treatment were harvested with a needle. Briefly, hyphae were immersed in a solution containing 0.2 M Tris-HCl pH 7.4, 1 M sodium succinate hexa-hydrate, 1 mg ml^-1^ nitro blue tetrazolium (NBT) and 5 mM MgCl_2_. The SDH, present in viable fungal hyphae, reacts with NBT and was reduced to a dark blue-violet formazan compound. The hyphae were washed with 1 ml of physiological water (0.9 % (w/v)). A second staining, to obtain better contrast, was performed with fuchsin acid (0.1% (v/v)). The samples were then cleaned in a solution of lactoglycerol. Pictures of the stained hyphae were taken in the brightfield channel by stereomicroscopy. A Nikon SMZ1270 stereomicroscope (Nikon, Japan) equipped with a Nikon DS-Qi2 monochrome microscope camera and a DS-F 2.5 F-mount Adapter 2.5× was used. The stereomicroscope was used with an ED Plan 2×/WF objective (Nikon, Switzerland) and an OCC illuminator allowing the image capture in the brightfield channel at an exposure time of 40 ms. The images were processed using Fiji^85^ by measuring, by threshold settings, the total surface of the hyphae and the surface corresponding to potential SDH activity. The SDH activity was calculated by the following formula: relative dark area 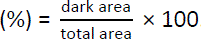. The quantification was performed from the biological material of minimum 3 plates (biological replicates) where minimum 1 hyphae have been harvested by plate (technical replicate).

### Impacts of BSMs on AM fungal hyphae

The impact of lipopeptides of *B. velezensis* (i.e. surfactin, iturin and fengycin) was tested on hyphae integrity. The potential permeabilization of the AM fungus membrane was quantified using the fluorescent intercalating propidium iodide (PI) that is not internalized by healthy cells. Four hyphae per Petri plate developing at the surface of the HC^Solid+Ri^ medium were exposed to one drop of 1 µl of 2 µM, 20 µM or 50 µM of iturin, surfactin or fengycin and incubated at room temperature for 30 minutes. The PI solution at 50 µg ml^−1^ was then applied on each hyphae for 15 min at room temperature in the dark. The stained hyphae were visualized by fluorescence microscopy (Nikon Ti2-E) with appropriate filters (TexasRed HC BrightLine Basic Filter). A control consisting of hyphae treated with PBS solution (Control) and 1% Triton X-100 with 2% formaldehyde (Killed) were considered. The fluorescence corresponding to PI was quantified using NIS-Element AR software (Nikon, Japan). The membrane permeabilization was assessed by threshold settings to obtain the region of interest (ROI) within the images taken in the bright field channel corresponding to hyphae area. Within this ROI, the mean of red intensity (MeanRed), equivalent at the arithmetic mean of pixel intensities, was quantified in the images taken in the red channel. The experiment was performed in 3 biological replicates (3 Petri plates) per treatment.

### Bacterial CFU counting *in vitro*

The colonization density of *B. velezensis* GA1 GFP on *D. carota* transformed roots (HC^solid+Dc^), *R. irregularis* hyphae (HC^solid+Ri^) and *S. tuberosum* roots (HC^solid+St^) was quantified. Bacterial inoculation was performed as previously described in the section “Root and hyphae colonization by *B. velezensis in vitro* “.

The colonization density of roots and hyphae as well as spores produced by the bacteria was evaluated 3, 7 and 14 dpi of *B. velezensis* for AM fungal hyphae and roots of *D. carota*, while for *S. tuberosum* roots, it was performed 14 dpi. Colonization was evaluated on 2 cm of roots or AM fungal hyphae proximal of the inoculation drop area. Bacterial cells were detached from roots and hyphae by vortexing for 5 min in a solution of physiological water supplemented with 0.1% (vol/vol) Tween 80 and 6 glass beads. To evaluate the number of bacterial spores, half of each solution was incubated at 80°C during 25 minutes to kill all vegetative cells. The colonies were counted in performing serial dilutions plated onto LB medium solidified with 14 g l^−1^ of agar. Plates were incubated for 12 h at 30°C. The quantification was performed from the biological material of minimum 4 replicates (i.e. plates) divided into 3 sample for each treatment. The results were expressed following the available area provided by each host depending of the sampling length (2cm) and of the average diameter of each host (Supplementary data Fig. 4).

### *B. velezensis* metabolite production along AM fungal hyphae

Metabolites produced by *B. velezensis* GA1 tagged GFP growing in contact with hyphae were extracted and analysed by LC-MS. As previously described in the section “Root and hyphae colonization by *B. velezensis in vitro*”, *B. velezensis* was inoculated on the surface of hyphae in the HC (HC^Solid+Ri^). After 9 days, a plug of agarose gel (0.5 cm × 2.5 cm) containing *R. irregularis* hyphae and the bacterium was sampled. BSMs were extracted via the application of 400 µl of acetonitrile (ACN) (75 %, vol/vol) during 30 min and then filtered with hydrophilic PTFE syringe filters ROCC S.A. (0.22 µm pore size) before UPLC-qTOF MS analysis.

Briefly, the extracts were analyses using a Agilent 1290 Infinity II apparatus coupled with a diode array detector and mass detector (Jet Stream ESI-Q-TOF 6530) with the parameters: capillary voltage: 3.5kV; nebulizer pressure: 35psi; drying gas: 8l/min; drying gas temperature: 300°C; flow rate of sheath gas: 11l/min; sheath gas temperature: 350°C; fragmentor voltage: 175V; skimmer voltage: 65V; octopole RF: 750V. Accurate mass spectra were recorded in positive mode in the range of m/z =100–1700. Metabolites (injection volume, 10 µl) were separated on C18 Acquity UPLC BEH column (2.1 × 50mm × 1.7μm; Waters, milford, MA, USA) and the following solvent system: acetonitrile and distilled water (both supplemented with 0.1% (vol/vol) formic acid), a flow rate of 0.6 ml min^−1^ and gradient over 20 min (programme: initial 10% (vol/vol) acetonitrile during 1 min before increased to 100% (vol/vol) over 20 min, held at 100% (vol/vol) for 3.5 min).

The identified BSMs were quantified by their peak area values using MassHunter v10.0 workstation. The Clips were quantified based on their retention times and masses compared with pure molecule standards.

The relative proportion of the detected CLiPs was then calculated by following formulas:

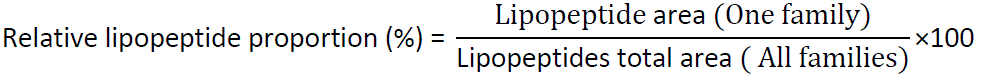

### *B. velezensis* metabolite production on *R. irregularis* exudates

The metabolite production of *B. velezensis* grown on *R. irregularis* exudates was analysed by LC-MS. To observe the effect of *R. irregularis* exudates on the BSMs production of GA1, the ERM of *R. irregularis* was cultivated on MSR^min-1/2^ liquid medium (HC^Liquid+Ri^). After 4 weeks of *R. irregularis* growth, the liquid medium was collected and hyphal exudates of 5 different plates pooled. At least 15 hyphal exudate solutions from 15 individual plates were grouped in 3 distinct solutions. Harvested exudates were freeze-dried (Freeze-dryer Alpha 3-4 LSCbasic brand Christ), resuspended and concentrated 5 times with a solution containing MOPS buffer (10.5 g l^-1^) and NH_4_SO_4_ (1 g l^-1^). The exudates concentrated 5 times were sterilized with CA syringe filters ROCC S.A. (0.22 µm pore size).

Then, the continuous growth kinetics of *B. velezensis* in AM fungal exudates was followed on microplates 96 wells. First, a pre-culture of *B. velezensis* was done at 30°C in liquid RE medium^48^ under agitation (180 rpm) overnight. The preculture was washed twice in physiological water (Na Cl 0.9 % w/v). The bacterial suspension was adjusted to obtain an OD_600_ of 0.05 into 96-well microtiter plates to apply final volume of 200 µl per well. The growth of *B. velezensis* was evaluated in 5 times concentrated exudates of *R. irregularis*. The growth kinetics of *B. velezensis* (OD_600_) was followed every hour during 36 h on a Tecan Spark automatic plate reader (Tecan Group Ltd, Männedorf, Switzerland) with continuous shaking at 30°C. To study the effect of hyphal exudates on the GA1 metabolome, 3 wells of each condition were pooled, filtered (0.22 µm) and then analysed by LC-MS as previously describe in the section:“ *B. velezensis* metabolite production along AM fungal hyphae.”. At least 3 distinct concentrated hyphal solutions (biological replicates) were inoculated with minimum 2 distinct *B. velezensis* pre-culture and cultured in 3 wells pooled after GA1 development (technical replicates) to perform LC-MS analysis.

### *B. velezensis* metabolite production on carbohydrate compounds present in *R. irregularis* exudates

The influence of carbohydrate compounds in the hyphal exudates of the AM fungus, often reported in the literature, was evaluated on BSMs production of GA1 WT. The carbon sources of AM fungus (10 mM of fructose, glucose, inositol, citric acid, 5 mM trehalose and 15 mM succinic acid) exudates were solubilized in M9 minimal salts medium (KH_2_PO_4_ 3 g L^-1^; NaCl 0.5 g L^-1^; Na_2_HPO_4_ 6.78 g L^-1^; NH_4_Cl 1 g L^-1^) supplemented with 2 mM MgSO_4_, 0.1 mM CaCl_2_, 10 μM FeSO_4_ and tamponed at pH 6.8 with MOPS (10.5 g l^−1^) forming the so-called AMF-EMM for Arbuscular Mycorrhizal Fungi Exudate Mimicking Medium. The metabolite production of GA1 was followed as previously described in the section “*B. velezensis* metabolite production along AM fungal hyphae”. The experiment was performed in 3 biological replicates (3 precultures of *B. velezensis*) with minimum 4 technical replicates for each biological replicate.

### *B. velezensis* antimicrobial activity assays on hyphal exudate

Antimicrobial activity of GA1 wild type (WT) or GA1 mutants’ cell-free supernatants (CFS) produced on hyphal exudates of *R. irregularis* was tested against *Trichoderma harzianum* Rifai MUCL 29707 and *Collimonas fungivorans* LMG 21973. For this, GA1 WT and mutants (Supplementary Table 1) were cultivated on exudates of *R. irregularis* as describe in the “B*. velezensis* metabolite production on *R. irregularis* exudates.”. The CFS of these cultures were obtained by centrifugation of the bacterial culture at 10000 rpm and sterilization with a PTFE syringe filters ROCC S.A. (0.22 µm pore size).

To observe the effects of GA1 WT or GA1 mutant’s CFS on the growth of *T. harzianum.*, fungal spores were harvested from 10 days old PDA cultures and suspended to a concentration of 2×10^5^ CFU mL^−1^ with physiological water supplemented with 0.1% (vol/vol) Tween 80. Ten µl of fungal spores were resuspended into 96-well microtiter plates with final volume of 200 µL adjusted with PDB medium per well supplemented with 20% (vol/vol) of GA1 WT or GA1 mutant CFS. The control was complemented with physiological water supplemented with 0.1% (vol/vol) Tween 80. The growth kinetics of *T. harzianum* (OD_600_) was followed every hour during 36 h with a Tecan Spark automatic plate reader (Tecan Group Ltd, Männedorf, Switzerland) with continuous shaking at 26°C. Three independent assays each involving minimum 4 technical replicates were performed.

To evaluated the impact of CFS on the growth of *C. fungivorans,* a bacterial suspension of this bacterium was prepared by centrifuging the culture overnight, washing the cells twice, and resuspending them in physiological water. The bacterial suspension was adjusted to obtain an OD_600_ of 0.05 into 96-well microtiter plates to apply final volume of 200 µl per well-adjusted with TSA medium (Sigma-Aldrich, India) supplemented with 20% (vol/vol) of GA1 WT or GA1 mutant CFS. The control was complemented with physiological water. The growth kinetics of *C. fungivorans* (OD_600_) was followed every hour during 24 h on a Tecan Spark automatic plate reader (Tecan Group Ltd, Männedorf, Switzerland) with continuous shaking at 26°C. Three independent assays each involving 6 technical replicates were performed.

### Translocation of surfactin via the AM fungal network

The transport of surfactin across the fungal network was assessed using the tri-compartment Petri plate system, as described in the “Velocity of Cytoplasmic Flow” section. To investigate the effectiveness of AM fungal hyphae in transporting surfactin, we applied 10 ml of a 20 µM surfactin solution to one of the hyphal compartments, covering the entire compartment (referred to as HC^solid+Ri+srf^). To determine whether surfactin was translocated through the ERM, we conducted a control experiment in which the *R. irregularis* network was cut before the surfactin application along the entire length of the compartment and on 0.5 cm of width, creating an air gap in the non-inoculated compartment. Subsequently, we collected the medium from both the treated and non-treated hyphal compartments. We then performed an extraction and LC-QTOF MS analysis of the surfactin contents of each hyphal compartment, following the procedures outlined in the “*B. velezensis* Metabolite Production Along AM Fungal Hyphae” section. This experiment was carried out in 3 biological replicates, with 3 plates for each condition.

### Experimental design of greenhouse trial for *B. velezensis* CMN colonization

Two 500 ml pots, designated as Pot 1 and Pot 2, were connect by a 4.5 cm diameter pipe made of HDPE (High-density polyethylene) (Fig. 2D). Both ends of the tube were sealed by nylon mesh of 50 µm porosity to prevent the passage of plant roots from one pot to the other. Each pot contained a sterile mixture of sand/vermiculite/loam (45%:45%:10%, v/v/v). One 20-day-old *S. tuberosum* plantlet produced *in vitro* was planted in each pot. Prior to the planting, 8 plants in pot 1 were inoculated with 10 g of *R. irregularis* inoculum (+ *R.i.*). The inoculum was obtained by associating isolated spores from an *in vitro* culture of the AM fungus with maize plants (*Zea mays* L. cultivar ES Ballade) grown in sterilized lava stone (DCM, Belgium) for 4 months. The roots and the rhizospheric attached from these plants were homogenized and used as *R. irregularis* inoculum. Additionally, 8 other plants of pots 1 were inoculated with 10 g of sterilized (121°C for 2h) *R. irregularis* inoculum (− *R.i.*). The plants were grown in greenhouse at 25 °C with a RH of 75% and 16/8 h (day/night) PPF of 120 mmol s^−1^ m^−2^. The plants were watered once a week with distilled water and fertilized once a week with Hoagland nutrient solution impoverished in phosphorus (Hoagland^-P^) ^87^. After 3 month of growth, each plant in pot 1 pot was watered with 50 ml of a bacterial suspension of *B. velezensis* GA1 GFP. This suspension was prepared by centrifuging a culture overnight (RE medium, 26°C), washing the cells twice, and resuspending them in Hoagland^-P^ to a density of 5 × 10^8^ CFU ml^−1^. The 16 uninoculated plants in pots 2 were treated with 50 ml of Hoagland^-P^ solution without any microorganisms. The colonization of *R. irregularis* and *B. velezensis* in roots of plants in pots 1 and in pots 2 was quantified 20 days after the inoculation of *B. velezensis* as described in section “Mycorrhizal root colonization under greenhouse trials” and “Bacterial root colonization under greenhouse trials”.

### Effect of *B. velezensis* colonization on AM fungal population under greenhouse trial

The potential influence of the bacterial colonization on the *R. irregularis* population was evaluated under greenhouse condition. Briefly, 16 pots were inoculated with 5 g of *R. irregularis* inoculum as previously described in the section “Experimental design of greenhouse trial for *B. velezensis* CMN colonization.”. At the same time, a sterilized seed of tomato cv Money Makers was sown in each pot. The plants were grown in greenhouse at 25 °C with a RH of 75% and 16/8 h (day/night) photoperiod under a PPF of 120 mmol s^−1^ m^−2^. The plants were watered once a week with distilled water and fertilized once a week with Hoagland^-P^. After 5 month, to observe the effect of *B. velezensis* on *R. irregularis* population, 8 pots containing plants were watered with 50 ml of a bacterial suspension of GA1 GFP as previously describe (*B.v.*+ *R.i.*). The remaining pots were watered with 50 ml of Hoagland^-P^ solution without *B. velezensis* (*R.i.*). The colonization of *R. irregularis* and *B. velezensis* in roots of plants for each condition was quantified before the bacterial inoculation and 7 days after the inoculation of *B. velezensis* as described in section “Mycorrhizal root colonization under greenhouse trials” and “Bacterial root colonization under greenhouse trials”.

### ISR induction of tomato plants under greenhouse condition

Sterilized Ailsa Criag seeds were transferred to 1 L pots containing a sterile substrate of sand/vermiculite/loam (45%:45%:10%,v/v/v). For each treatment, 16 pots were used. Thus, 16 pots received 5 g of *R. irregularis* inoculum containing AM fungal-colonized roots and spores (*R.i.*) and prepared as previously describe in the section “ Experimental design of greenhouse trial for *B. velezensis* CMN colonization”. The inoculation of tomato plants with *R. irregularis* was done by mixing the substrate with the AM fungal inoculum prior to sowing.

Likewise, 16 plants were watered with 10 ml of a bacterial suspension (*B.v.*) as described previously in “ Experimental design of greenhouse trial for *B. velezensis* CMN colonization”. Additionally, 16 pots were co-inoculated with both the bacteria and the AM fungus (*R.i.* +*B.v.*) as described for treatment with one microorganism. A second bacterial application was performed after 12 weeks of plant growth by watering the plants with 10 ml of a bacterial suspension at the same concentration. A control group composed of 16 non-inoculated plants was treated with Hoagland^-P^ solution without the bacterial suspension and receiving the same amount (5 g) of AM fungal sterilize inoculum substrate.

The plants were maintained in the greenhouse and watered once a week with distilled water non sterilized. They were also fertilized once a week with Hoagland^-P^. The pots were arranged in a fully randomized design. The plants were grown in the greenhouse at 25 °C with a RH of 75% and 16/8 h (day/night) photoperiod under a PPF of 120 mmol s^−1^ m^−2^ for 3 months.

At week 13, the plants were infected with *B. cinerea* MUCL 43839. Prior to infection, the size of the plants was measured to assess the impact of the combined treatment on plant growth compared to the control plants (n=16/treatment). *B. cinerea* was routinely cultured on potato dextrose agar at 26°C from a spore suspension stored at –80°C. *B. cinerea* spores were collected from 15-days old cultures in physiological water containing 0.01% Tween 20. The spore suspension was then filtered, quantified and adjusted to a concentration of 5 × 10^6^ spores ml^−1^. Infection was performed by applying 10 μl droplets of the spore suspension onto 5 leaflets of the third, fifth and sixth leaves (15 infected leaflets/plant). The disease severity of *B. cinerea* on tomato leaves was assessed by measuring the lesion area using ImageJ 14 dpi (n=8/treatment).

The colonization of *R. irregularis* and *B. velezensis* in the combined treatment (*B.v.* + *R.i.*) was quantified 14 and 21 dpi after the second GA1 treatment (n=8/times) as described in section “Mycorrhizal root colonization under greenhouse trials” and “Bacterial root colonization under greenhouse trials”.

### Bacterial root colonization under greenhouse trials

To determine *B. velezensis* colonization, bacterial cells were detached from 1 g of roots and attached rhizospheric substrate samples by vortexing for 5 min in a solution of physiological water supplemented with 0.1% (vol/vol) Tween 80 and 6 glass beads. The colonies were counted by performing serial dilutions plated onto LB medium supplemented with chloramphenicol 5 µg l-1. Plates were incubated for 12h at 30°C, and the results were expressed as the number of CFU of *B.v.* g-1 of sample.

### Mycorrhizal root colonization under greenhouse trials

To determine the mycorrhizal status of plant, roots with attached rhizospheric substrate were sampled to quantify the presence of *R. irregularis* by qPCR. Frozen roots were crushed in liquid nitrogen, and DNA was extracted from a 500 mg crushed root sample using GenEluteTM Soil DNA Isolation kit (Sigma-Aldrich, Canada) with slight modifications. Prior to extraction, a lysis step was performed using DNAzol™ Reagent under shaking and homogenization with FastPrep-24™(MP Biomedical’s, Germany). The concentration and purity of the extracted DNA were assessed with the UV-vis spectrophotometer NanoDrop 2000 (Thermo scientific). qPCR was performed in a ABI StepOne™ qPCR apparatus (Applied Biosystems) using the kit Luna® Universal qPCR Master Mix Kit (New England Biolabs, Ipswich, MA, United States). PCRs were conducted in a total volume of 20 µl containing 10 μL of Luna Universal qPCR Mix, 0.5 μl of each primer (10 µM) and 5 µl of DNA. The qPCR protocol consisted of an initial denaturation step at 95°C (1 min), followed by 40 cycles of denaturation (95°C, 15 s) and extension (60°C, 30 s). Melting curves were also generated from 65 to 95°C with an increase rate of 0.5°C/5s to evaluate the specificity of the amplified products. The strain-specific primer pair for *R. irregularis* MUCL41833 targeting the mtLSU region was used: forward 5’-AAGTCCTCTAGGTCGTAGCA-3’ and reverse 5’-ACAGGTATTTATCAAATCCTTCCC-3’. The resulting concentrations were expressed as mg of R.i. g-1 of sample. The quantification of *R. irregularis* was performed based on the standard calibration curves. To prepare standards for the qPCR experiments, DNA extracted from *R. irregularis* spores and mycelium grown in vitro were used. Spores/mycelium of 3 month old cultures were extracted from HC following solubilization of the phytagel by citrate-buffer. Five mg of spores/mycelium was quantified and lysed with DNAzoleTM. A serial 4-fold dilutions of the lysed extraction with DNAzole (5×10−1–10−4 mg μl−1) was performed prior to the DNA extraction, following the same procedure described before for roots.

### Statistical analysis

Data were processed using GraphPad Prism 9.2.0 software to perform statistical analysis with a Student’s paired *t* tests. For multiple comparisons, results were analysed by one-way analysis of variance (ANOVA) with a general linear model procedure of data from several (mostly three biological replicates) independent experiments and Tukey’s honestly significant difference (HSD) tests. The impact of factors (experiments, treatments, biological materials and dpi) on variability were considered in the statistical general analyses to analyse repeated measure experiments. Differences were considered statistically significant at p < 0.05.

## Data availability

Source data for all relevant figures are provided with this paper.

## Acknowledgements

We gratefully acknowledge Andrew Zicler for technical help with the establishment of the experimental set-up in microscopy. We thank Jos Raaijmakers (Netherlands Institute for Ecology, Wageningen) and Monica Höfte (Ghent University) for their critical reading of the manuscript.

## Authors Contribution

A.A., S.D. and M.O. conceived, designed, and coordinated the project. A.A., L.A.P, S.L., C.H., F.B., S.S., and A.A.A. generated materials, performed experiments, and/or analyzed data. A.A. and M.O. wrote the manuscript. A.A, S.D., M.C.S. and M.O. revised the manuscript.

