## Supplementary for "Deciphering the biology and chemistry of the mutualistic partnership between *Bacillus velezensis* and the arbuscular mycorrhizal fungus *Rhizophagus irregularis*"

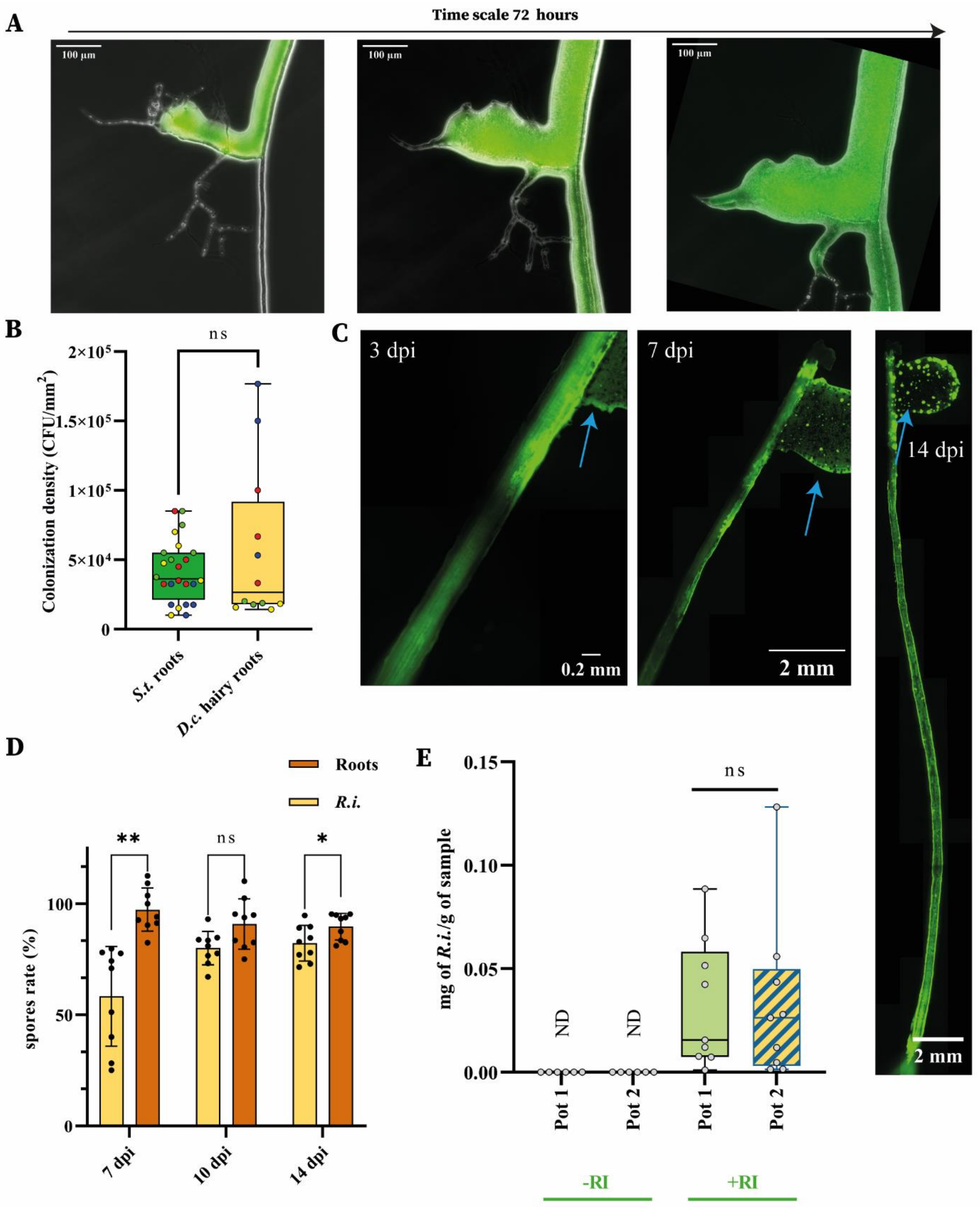


**Supplementary Data Fig. 1:** ***B. velezensis*** **colonization along roots and AM fungal hyphae. A**, Microscopic pictures of *B. velezensis* colonization along hyphae of a 3 month-old culture of *R. irregularis* during a time scale of 72 hours showing the development of a biofilm and colonization of new sections of the hyphae. **B**, Colonization density of 3 months old cultures of Solanum tuberosum (*S.t.)* roots or transformed roots of Daucus carota (D.c.) 14 dpi of *B. velezensis.* The boxes encompass the 1st and 3rd quartiles, the whiskers extend to the minimum and maximum points, and the midline indicates the median. The individual points represent 4 biological replicates (different colours) and 3 at 6 technical replicates (same colours). 12≤ n ≤ 24; student's t test (α = 0.05): ns = not significant**. C**,

*B. velezensis* colonization along a D. carota. transformed root. Microscopic composite pictures were taken in epifluorescence. The blue arrow show the inoculation drop area on the root. The speed of colonization was quantified by the measurement of the distance travelled by *B. velezensis* (green colour) from the inoculation drop (Blue arrow) divided by the day post inoculation (dpi). **D**, Rate of *B. velezensis* spores formed during its colonization along hyphae of 3 month old *R .irregularis* and hairy roots of 3 month old *D. carota*, 7, 10 and 14 dpi. Bars represent the means ± SD of 9 biological replicates. n =9; ns = not significant; Mixed-effects model and Šídák's multiple comparisons test ($p < 0.05$): * $0.01 < p < 0.05$; ** $0.001 < p < 0.01$. **E**, Fungal population of *R. irregularis* quantified by qPCR reported by gram of sample (mg of *R.i.* /g of sample). All the plants in pot 1 associated (+R.i.) or not (-R.i.) with *R. irregularis* were inoculated with *B. velezensis*. Plants in pot 2 were not inoculated with any microorganisms. The individual points represent 9 biological replicates (9 systems). n =9; one-way analysis of variance (ANOVA) and Tukey's HSD test (Honestly significantly different, α = 0.05): Groups with different letters differed significantly from each other at an α of 0.05.ND= no detected.

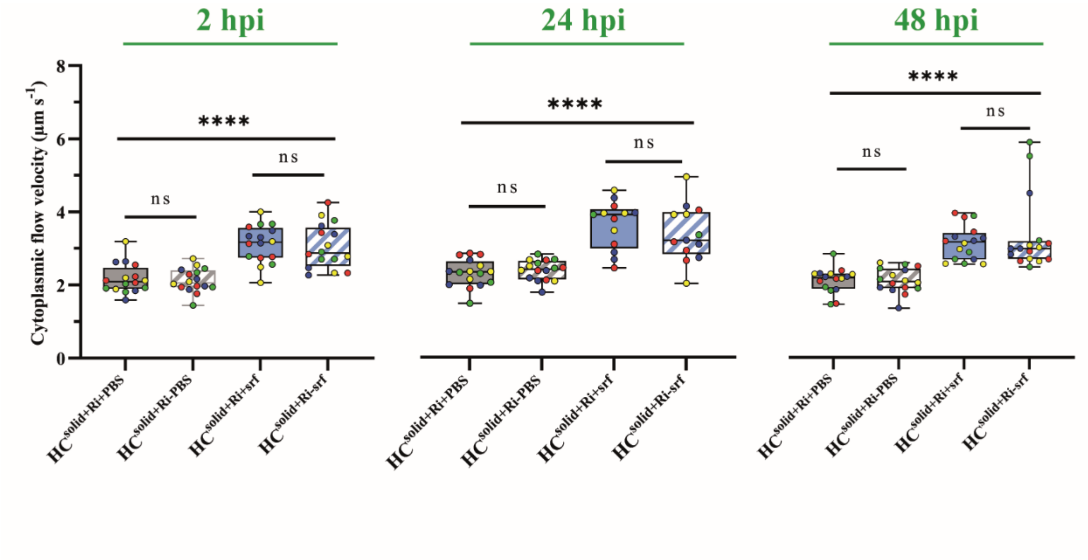


**Supplementary Data Fig. 2:Dynamics of *R. irregularis* cytoplasmic flow velocity in presence of surfactin**. The velocity of the cytoplasmic flow in the compartment treated with surfactin (HCSolid+Ri+srf , blue) or PBS (HC Solid+Ri+PBS, gray) and in the untreated compartment following the treated compartment either with surfactin (HC Solid+Ri-srf, blue stripes) or with PBS (HC Solid+Ri-PBS, gray stripes) after 2, 24 and 48 hours of treatment. The boxes encompass the 1st and 3rd quartiles, the whiskers extend to the minimum and maximum points, and the midline indicates the median. The individual points represent 4 biological replicates (different colours) and 4 technical replicates (same colours). n=16; one-way analysis of variance (ANOVA) and Tukey's HSD test ($\alpha$ = 0.05): ns = not significant; **** $p < 0.0001$.

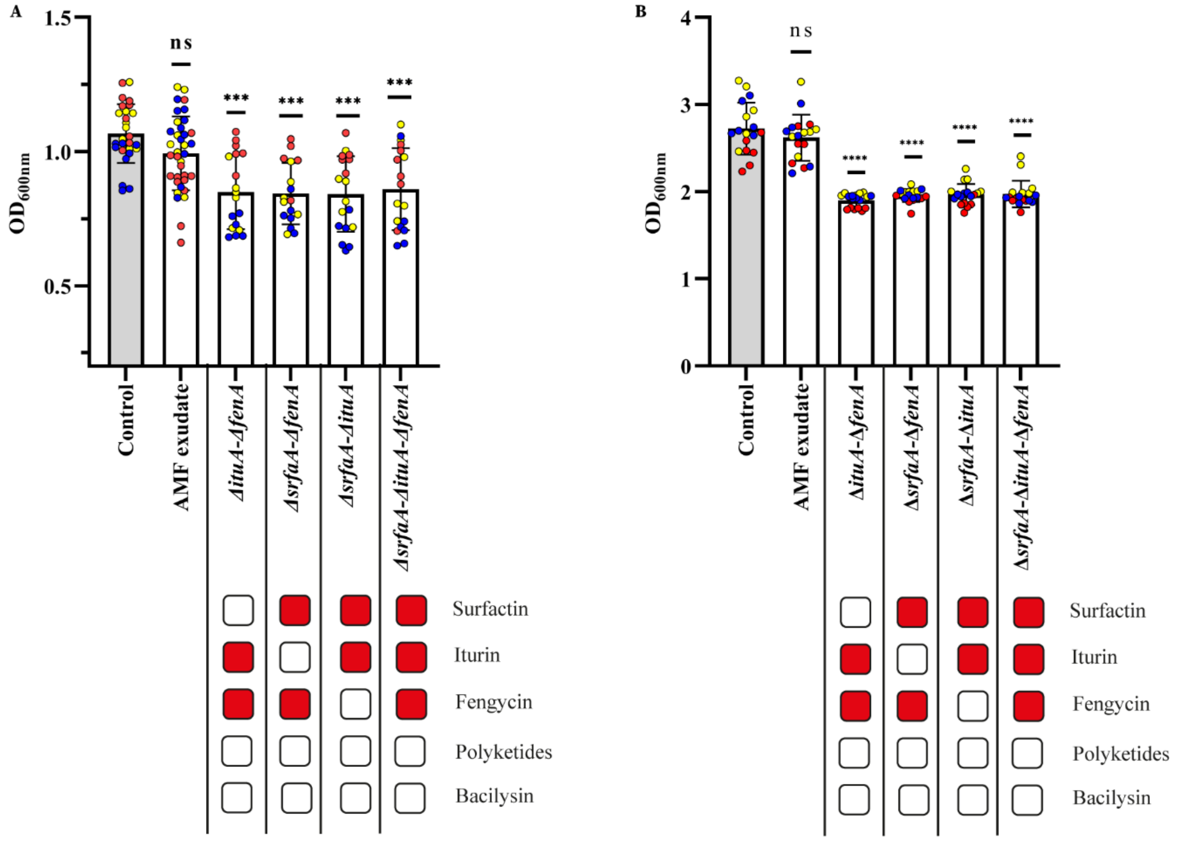


**Supplementary Data Fig. 3:BSMs produced by *B. velezensis* on AM fungal exudates allowing an antagonism activity against *Trichoderma harzianum* and *Collimonas fungivorans*. A**, Effect of *B. velezensis* GA1 wild type or mutant cell-free supernatants (CFS), produced on exudate of *R. irregularis*, on the growth of *Trichoderma harzianum* Rifai MUCL 29707. The optical density (OD600) of the mycoparasite fungus was quantified after 36 h of growth in presence or in absence (control) of CFS. Metabolites not produced by the different mutants are illustrated with red boxes in the table above. Bars represent the means ± SD of 3 biological replicates (different colours) and 4 to 9 technical replicates (same colours). $16 \leq n \leq 27$, one-way analysis of variance (ANOVA) and Tukey's HSD test ($\alpha = 0.05$): ns = not significant; **** = $p < 0.0001$. **B,** Effect of *B. velezensis* GA1 wild type or mutant cell-free supernatants (CFS), produced on exudate of *R. irregularis.,* on the growth of *Collimonas fungivorans* LMG 21973. The optical density (OD600) of the bacteria was quantified after 36 h of growth in presence or in absence (control) of CFS. Metabolites not produced by the different mutants are illustrated with red boxes in the table above. Bars represent the means ± SD of 3 biological replicates (different colours) and 6 technical replicates (same colours). $16 \leq n \leq 27$, one-way analysis of variance (ANOVA) and Tukey's HSD test ($\alpha = 0.05$): ns = not significant.

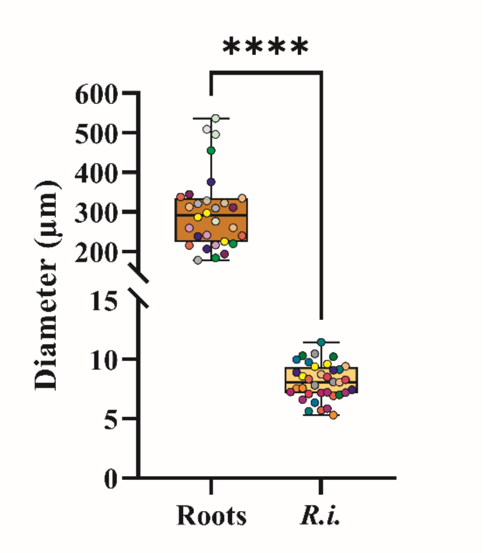


**Supplementary Data Fig. 4: Determination of the available surface provide by *D. carota* and *R. irregularis* for the bacterial colonization**. Diameter of D .carota roots or *R. irregularis* hyphae quantified by ImageJ Fiji. The boxes encompass the 1st and 3rd quartiles, the whiskers extend to the minimum and maximum points, and the midline indicates the median. The individual points represent 10 to 12 biological replicates (different colours) and 3 technical replicates (same colours). $30 \leq n \leq 36$. student's t test ($\alpha = 0.05$): **** = $p < 0.0001$

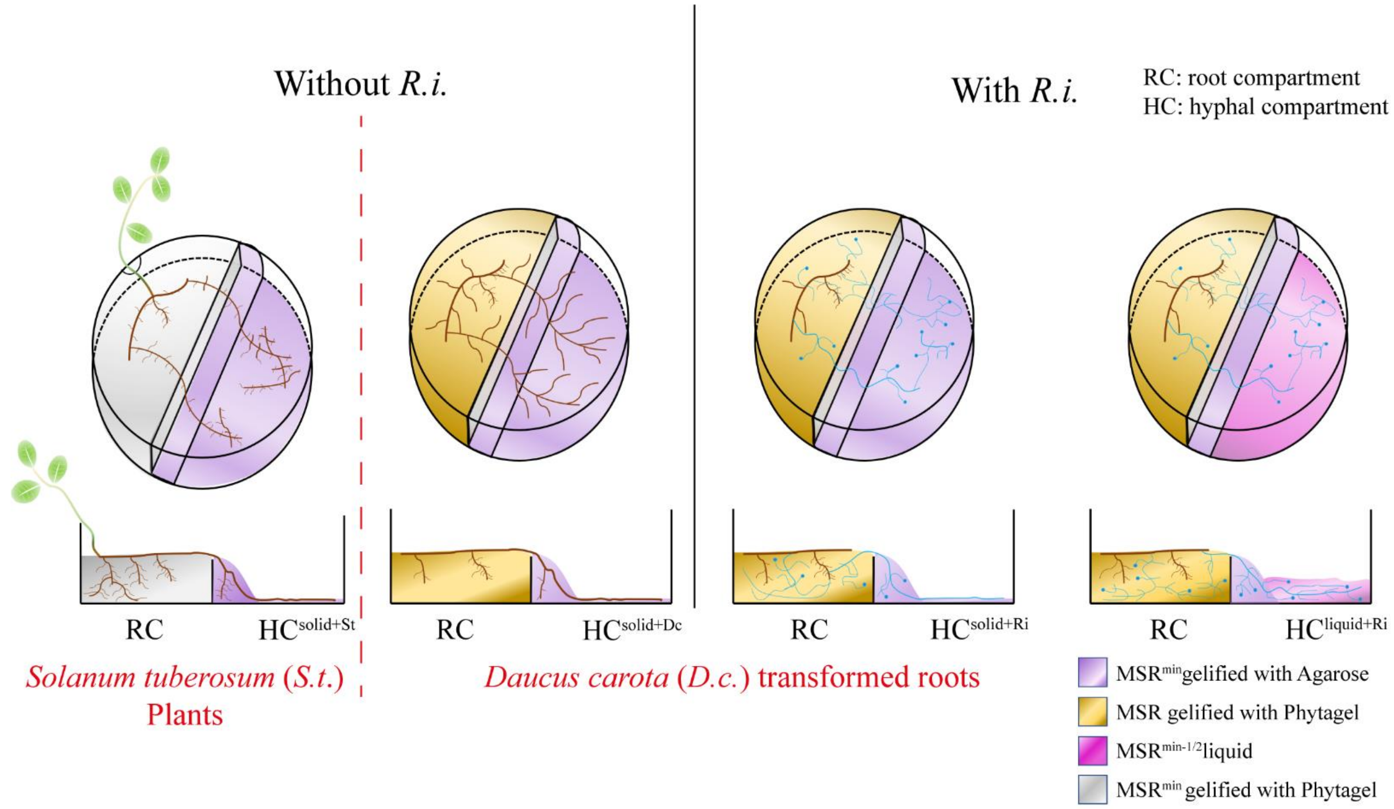


**Supplementary Figure 5: Schematic representation of experimental set-up**. Schematic view of systems consisted of bi-compartement Petri plates with a root compartment (RC) containing roots of *S. tuberosum* plants or roots of *D. carota* clone DC2 associated or not *to R. irregularis* and a hyphal compartment (HC) containing only roots of *S. tuberosum* (HCsolid+St) or of *D. carota* ( HCsolid+DC) or hyphae of *R. irregularis* (HCsolid+Ri and HCliquid+Ri). In the HC, roots or ERM were allowed to proliferate on solid (HCsolid+St, HCsolid+Dc and HCsolid+Ri) or in liquid (HCliquid+Ri) minimal media without any added carbon source. Brown, transformed roots of *D. carota* or roots *of S. tuberosum;* Blue, ERM of *R. irregularis*.

### Supplementary Table 1: Bacterial strains used in this study:

| | Relevant genotype, description (Genome bank number when sequenced) | Reference or sources |
|---|---|---|
| ***Bacillus velezensis*** | | |
| GA1 | Wild type (CP046386) | [118] |
| GA1 Δ*amyE*::cat+P*veg* -*gfpmut3* (called GA1 GFP) | GA1 disrupted of *amyE* gene; $Cm^R$ and harbouring a constitutive transcriptional fusion (P*veg*-*gfpmut3*) | [43] |
| GA1 Δ*srfaA* ::cat-Δ*fenA* ::phleo | GA1 deleted of *srfaA* and *fenA* genes; unable to produce surfactin and fengycin family | This study |
| GA1 Δ*srfaA* ::cat-Δ*ituA* ::phleo | GA1 deleted of *srfaA* and *ituA* genes; unable to produce surfactin and iturin family | This study |
| GA1 Δ*fenA* ::cat-Δ*ituA* ::phleo | GA1 deleted of *fenA* and *ituA* genes; unable to produce fengycin and iturin family | This study |
| GA1 Δ*fenA* ::cat-Δ*ituA* ::phleo-Δ*srfaA* ::kana | GA1 deleted of *fenA*, *ituA* and *srfaA* genes; unable to produce surfactin, fengycin and iturin family | This study |
| GA1 Δ*sfp* ::cat | GA1 deleted of *sfp* gene; unable to produce lipopeptides and polyketides | [42] |
| GA1 Δ*bacA* ::cat Δ*sfp* :;phleo | GA1 deleted of *bacA* and *sfp* genes; unable to produce bacilysin, lipopeptides and polyketides | This study |
| GA1 Δ*srfaA* ::cat | GA1 deleted of *srfaA* gene; unable to produce surfactin family | [43] |
| GA1 Δ*bacA* ::cat | GA1 deleted of bacA gene; unable to produce bacilysin | [42] |
